# PIGBOS-CLCC1 Interaction Shapes Cellular Calcium Dynamics and Energy Metabolism

**DOI:** 10.64898/2026.01.08.697870

**Authors:** Seemanti Aditya, Amal Kanti Bera

## Abstract

PIGBOS is a recently identified 54-amino acid microprotein localized to the mitochondrial outer membrane and implicated in the endoplasmic reticulum (ER) stress response. Here, we identify a previously unrecognized role for PIGBOS in cellular Ca^2+^ homeostasis. Manipulation of PIGBOS expression in HEK293T cells revealed that PIGBOS enhances Ca^2+^ signaling by promoting ER Ca^2+^ release through inositol 1,4,5-trisphosphate (IP_3_) receptors and subsequent mitochondrial Ca^2+^ uptake in response to histamine stimulation. In contrast, siRNA-mediated depletion or genetic ablation of PIGBOS markedly attenuated these responses. PIGBOS selectively facilitated Ca^2+^ transfer from the ER to mitochondria without affecting mitochondrial Ca^2+^ uptake from non-ER sources and also promoted store-operated Ca^2+^ entry. Functional analyses demonstrated that the interaction of PIGBOS with the ER-resident chloride channel CLCC1 via its C-terminal region is required for this activity. Network analysis predicted a direct association between PIGBOS and CLCC1, as well as indirect connections with core Ca^2+^ signaling components, including IP_3_ receptors, STIM1, Orai1, and SERCA, whose expression was altered upon modulation of PIGBOS abundance. Loss of PIGBOS impaired mitochondrial respiration, reduced ATP production, and increased reactive oxygen species. Together, these findings establish PIGBOS as a key regulator of ER-mitochondrial Ca^2+^ signaling that couples Ca^2+^ dynamics to mitochondrial bioenergetics and cellular stress responses.

**Significance Statement:** This study identifies PIGBOS, a mitochondrial microprotein, as a key regulator of cellular Ca^2+^ signaling through its interaction with the endoplasmic reticulum (ER) chloride channel CLCC1. By coordinating ER–mitochondrial Ca^2+^ transfer and modulating the expression of key Ca^2+^ signaling–associated proteins, PIGBOS integrates cellular Ca^2+^ dynamics with mitochondrial energy metabolism and stress responses. Given that perturbations in Ca^2+^ homeostasis profoundly influence cell survival and energy production, these findings reveal a new paradigm of inter-organelle communication. This work has broad implications for understanding the molecular basis of disorders such as neurodegeneration and cancer, where Ca^2+^ signaling and homeostasis are disrupted.

## Introduction

Calcium ion (Ca^2+^) regulates a broad spectrum of physiological processes, including muscle contraction, neurotransmitter release, fertilization, cell division, survival, and apoptosis (1–6). Cytosolic Ca^2+^ concentration is tightly maintained within the nanomolar-to-micromolar range through the coordinated actions of ion channels, receptors, transporters, and calcium-binding proteins (7, 8). The endoplasmic reticulum (ER) serves as the principal intracellular Ca^2+^ reservoir, releasing or sequestering Ca^2+^ in response to cellular cues (9). Activation of Gq-class G protein–coupled receptors increases inositol 1,4,5-trisphosphate (IP₃) production, which in turn activates IP₃ receptors (IP_3_R) to release ER Ca^2+^ ([Ca^2+^]_ER_) stores (10). The sarco/endoplasmic reticulum Ca²⁺-ATPase (SERCA) pump subsequently replenishes [Ca^2+^]_ER_, thereby maintaining steady-state levels essential for ER functions such as protein folding (11). Disruption of [Ca^2+^]_ER_ homeostasis impairs Ca^2+^-dependent chaperones, leading to protein misfolding and ER stress; if unresolved, prolonged ER stress can culminate in apoptosis (12, 13). Dysregulation of intracellular Ca^2+^ homeostasis has been implicated in neurodegenerative diseases, cardiovascular disorders, and cancer (14–17). The ER also forms specialized contact sites with mitochondria. These junctions, termed mitochondria-associated membranes (MAMs), facilitate Ca^2+^ transfer and lipid exchange between the two organelles (18, 19).

Recently, a 54-amino-acid microprotein, PIGB opposite strand 1 (PIGBOS), was identified on the mitochondrial outer membrane, where it interacts with the ER-localized chloride channel CLIC-like 1 (CLCC1) (20). Deletion of PIGBOS sensitized HEK293 cells to chemically induced ER stress, suggesting a role in ER homeostasis (20). Although PIGBOS does not influence the physical juxtaposition of the ER and mitochondria and is not a structural tethering component of MAMs, its interaction with CLCC1 (20) likely contributes to ER-mitochondria functional coupling. Notably, CLCC1 knockdown also induces ER stress, characterized by increased chaperone protein GRP78 expression and impaired protein folding (21). Moreover, amyotrophic lateral sclerosis (ALS) associated CLCC1 mutations disrupt ER luminal ion homeostasis (K^+^, Ca^2+^, Cl^-^), thereby exacerbating ER stress, possibly through ionic imbalance (22). However, the precise role of PIGBOS in ER function and the cellular stress response remains poorly understood. Given the involvement of CLCC1 in [Ca^2+^]_ER_ regulation and its interaction with PIGBOS, this complex may represent a critical modulator of Ca^2+^ signaling, particularly in mediating ER-to-mitochondria Ca^2+^ transfer.

In this study, we manipulated PIGBOS expression to investigate its role in Ca^2+^ signaling and mitochondrial function in HEK293T cells. Both overexpression and knockdown or knockout of PIGBOS significantly altered cellular Ca^2+^ signaling dynamics, especially the transfer of Ca^2+^ from the ER to mitochondria. These changes were accompanied by differential expression of multiple Ca^2+^ signaling-related genes and alterations in mitochondrial energy production. The interaction between PIGBOS and CLCC1 proved essential, as deletion of PIGBOS’s C-terminal CLCC1-interacting domain attenuated these effects. Notably, CLCC1 knockdown recapitulated the phenotypes observed upon PIGBOS depletion. Collectively, our findings identify PIGBOS as an integral component of the cellular Ca^2+^ signaling machinery, regulating Ca^2+^ homeostasis and mitochondrial metabolism through its association with CLCC1. Communication between organelles via the PIGBOS-CLCC1 axis governs cellular sensitivity to ER stress and, when dysregulated, may contribute to disease pathogenesis.

## Results

### PIGBOS regulates ER, cytosolic, and mitochondrial Ca^2+^ dynamics

To investigate the role of PIGBOS in cellular Ca^2+^ homeostasis, we expressed FLAG-tagged PIGBOS in HEK293T cells and confirmed its expression by western blotting (Fig. 1A). The mitochondrial localization of the overexpressed protein was verified by confocal microscopy (Fig. S1A). Ca^2+^ signaling was induced using histamine, which activates histamine receptors and triggers the generation of IP_3_. The resulting IP_3_ binds to IP_3_ receptors on the ER, leading to Ca^2+^ release from ER stores. The histamine-induced increase in cytosolic Ca^2+^ concentration ([Ca^2+^]_cyto_), measured using jGCaMP7s, was significantly higher in PIGBOS-overexpressing cells than in mock-transfected controls (Fig. 1B). One possible explanation for this enhanced response is an increased [Ca^2+^]_ER_ pool. To test this, we measured basal [Ca^2+^]_ER_ in unstimulated cells using GCEPIA1-SNAP_ER_, as described in the methods section. Steady-state [Ca^2+^]_ER_ was significantly higher in PIGBOS-overexpressing cells, indicating that PIGBOS can modulate basal [Ca^2+^]_ER_ content despite its predominant mitochondrial localization (Fig. 1C).

**Fig. 1.**
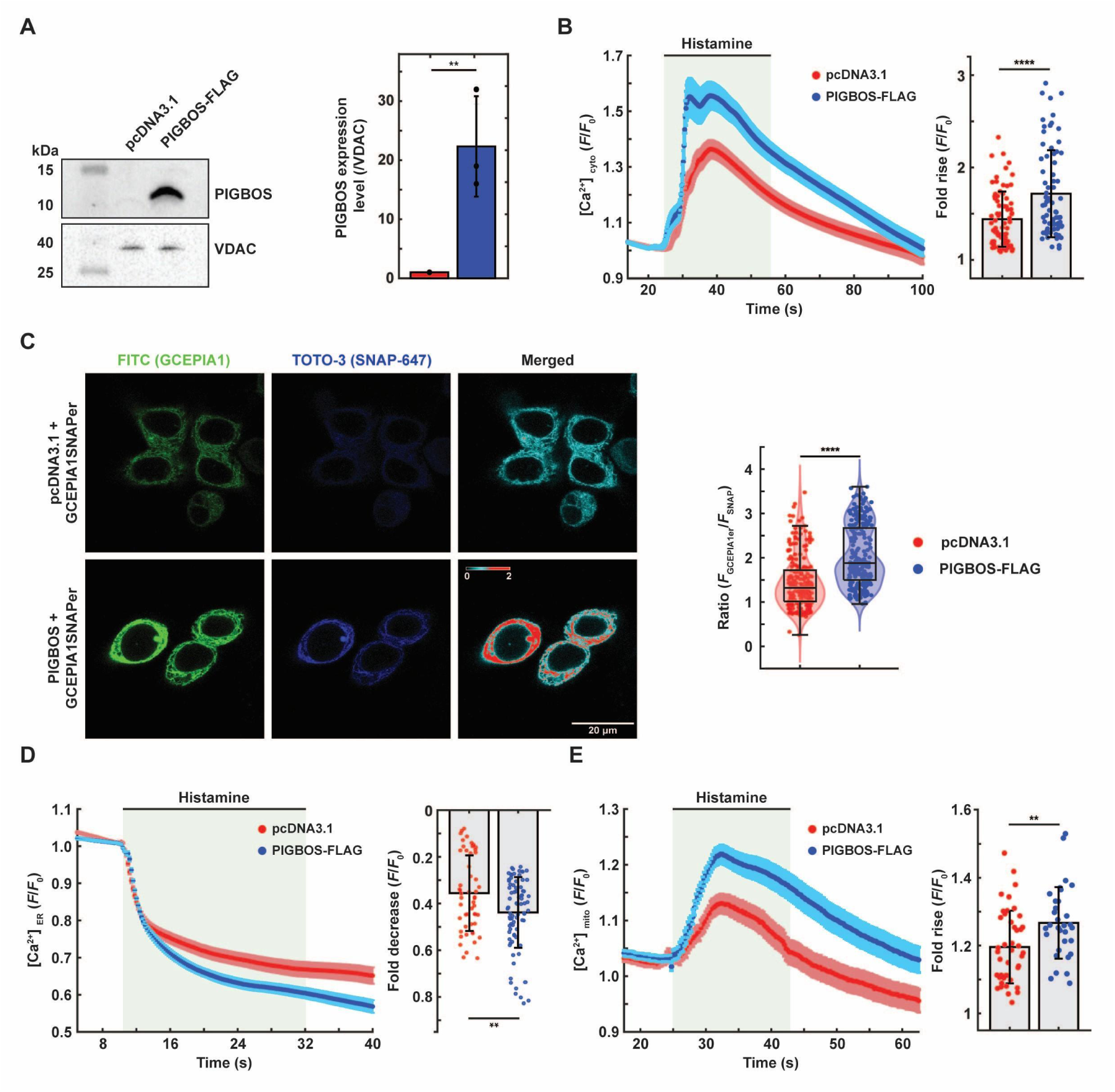
Effects of PIGBOS overexpression on Ca^2+^ dynamics. (A) Western blot confirms elevated PIGBOS protein levels in PIGBOS1-transfected HEK293T cells. PIGBOS expression, normalized to VDAC (loading control), is quantified from three independent experiments (bar graph). (B) PIGBOS overexpression enhances the histamine-induced rise in cytosolic Ca^2+^ (*n* = 75-76 cells) relative to mock-transfected controls. (C) Representative confocal images show elevated basal ER Ca^2+^ levels ([Ca^2+^]_ER_) in PIGBOS-overexpressing cells. Normalized GCEPIA1 intensity relative to SNAP signal (right; summarized from 230-260 cells). (D) Histamine-induced ER Ca^2+^ release is augmented in PIGBOS-overexpressing cells (bar graph; 52-81 cells). (E) PIGBOS overexpression increases histamine-stimulated mitochondrial Ca^2+^ uptake, measured with GCEPIA-2mt (bar graph; 43-48 cells). Data are mean ± SD. *P < 0.05, **P < 0.01, ***P < 0.001, ****P < 0.0001 (unpaired t test).

We next analyzed [Ca^2+^]_ER_ release following histamine stimulation. Direct measurement of [Ca^2+^]_ER_ with GCEPIA1-SNAP_ER_ showed that histamine-induced depletion was greater in magnitude in PIGBOS-overexpressing cells, compared with the control cells. Within 30 s of histamine application, fluorescence (F/F_0_) decreased by approximately 45% in PIGBOS-overexpressing cells versus ∼35% in controls, consistent with the enhanced [Ca^2+^]_cyto_ rise (Fig. 1D). A substantial fraction of the released Ca^2+^ is rapidly taken up by mitochondria at ER-mitochondria contact sites (MAMs) to restore the cellular Ca^2+^ homeostasis. Mitochondrial Ca^2+^ uptake, measured using GCEPIA1-2mt, indicated markedly increased levels in PIGBOS-overexpressing cells upon histamine stimulation (Fig. 1E). To verify that the FLAG epitope did not affect the observed results, the experiments were repeated using untagged PIGBOS, cloned into the IRES-mCherry vector (pICherryNeo). Comparable results were obtained with untagged PIGBOS, confirming that the findings were not influenced by the presence of the FLAG tag (Fig. S1B).

To address potential non-physiological effects of overexpression, we examined the effects after siRNA-mediated PIGBOS knockdown and CRISPR-Cas9 knockout (KO) in HEK293T cells. Efficient knockdown of PIGBOS was verified by western blotting. siRNA transfection reduced PIGBOS protein levels by approximately 70% (Fig. 2A). The generation of the KO line is described in the methods and summarized in Fig. S1C. Upon histamine stimulation, PIGBOS-deficient cells displayed the opposite phenotype of overexpressing cells: a smaller [Ca^2+^]_cyto_ transient (Fig. 2B), lower basal [Ca^2+^]_ER_ (Fig. 2C), reduced [Ca^2+^]_ER_ release (Fig. 2D), and diminished mitochondrial Ca^2+^ uptake (Fig. 2E). Similarly, PIGBOS-KO cells exhibited attenuated [Ca^2+^]_cyto_ elevations and reduced mitochondrial Ca²⁺ uptake following histamine stimulation (Fig. S2A-B).

**Figure 2.**
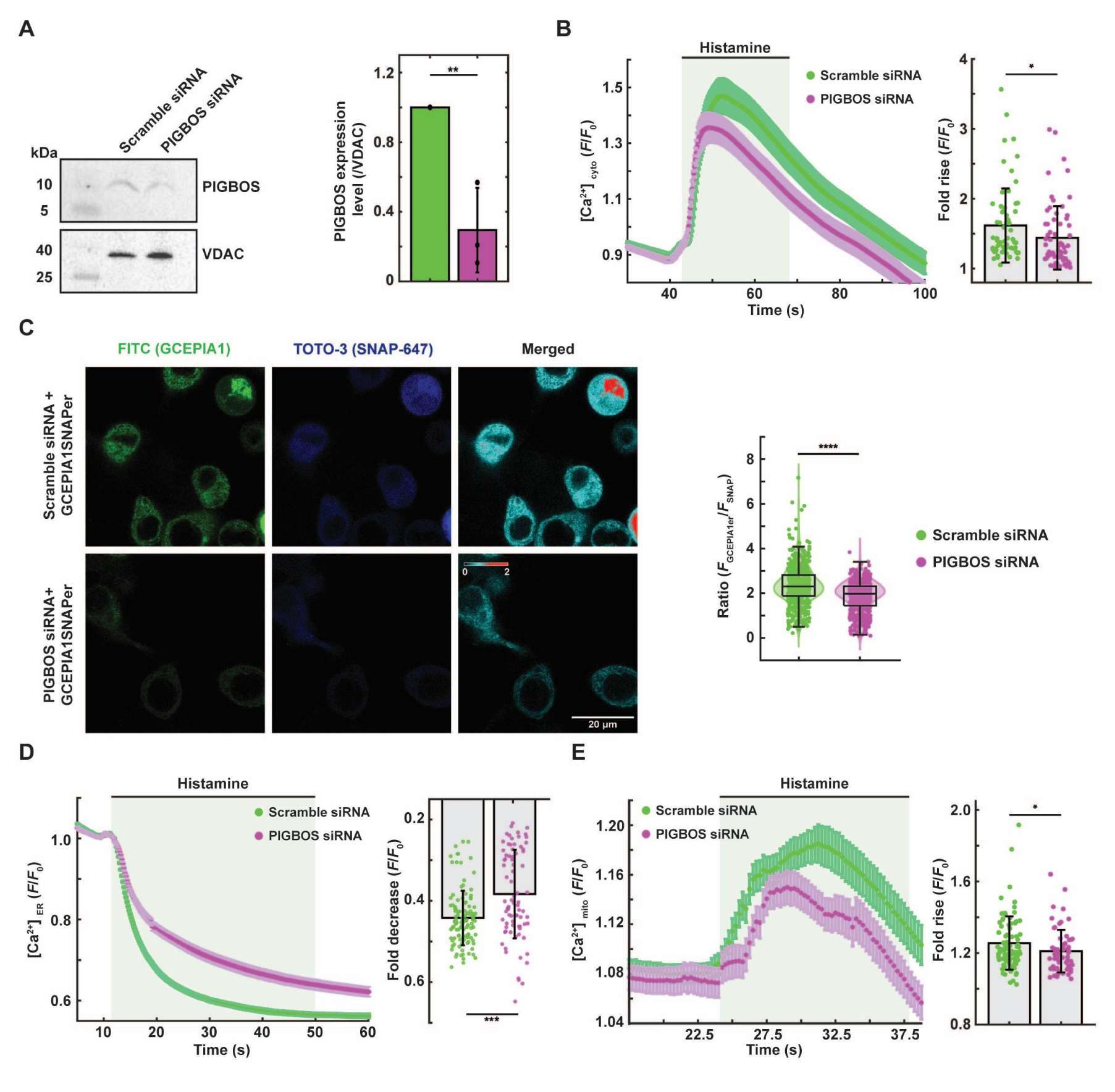
PIGBOS knockdown impairs cellular Ca^2+^ dynamics. (A) Western blot showing reduced PIGBOS expression following siRNA-mediated knockdown in HEK293T cells. VDAC was used as a loading control. Quantification from three independent experiments indicates an approximately 70% decrease in PIGBOS protein levels. (B) Knockdown of PIGBOS attenuates the cytosolic Ca^2+^ response to histamine stimulation. Bar graph shows the mean fluorescence ratio (F/F_0_) from 64-66 cells measured using jGCaMP7s. (C) Representative confocal images showing reduced basal ER Ca^2+^ levels in PIGBOS knockdown cells compared with scramble siRNA controls. The violin plot (right) summarizes data from 550-635 cells measured using G-CEPIA1-SNAP_ER_. (D) Histamine-induced ER Ca^2+^ release is diminished in PIGBOS knockdown cells (*n* = 80) relative to scramble controls (*n* = 114). (E) Mitochondrial Ca^2+^ uptake following histamine stimulation is reduced in PIGBOS knockdown cells (*n* = 63) compared with scramble controls (*n* = 86), measured using G-CEPIA1-2mt. Data are shown as mean ± SD. *P < 0.05, **P < 0.01, ***P < 0.001, ****P < 0.0001 (unpaired t test).

### PIGBOS selectively promotes ER-to-mitochondria Ca^2+^ transfer

Both overexpression and depletion experiments identified PIGBOS as a positive regulator of mitochondrial Ca^2+^ uptake. To assess whether this effect depends on ER-derived Ca^2+^, cells were permeabilized with digitonin in Ca^2+^-free buffer, and ER stores were depleted with thapsigargin (TG) in the presence of EGTA (Fig. 3A). Subsequent reintroduction of extracellular Ca^2+^ triggered robust mitochondrial Ca^2+^ uptake in both control and PIGBOS-overexpressing cells, with no significant difference in amplitude (Fig. 3B). These results indicate that PIGBOS selectively facilitates mitochondrial Ca^2+^ uptake from ER-released, but not extracellular, Ca^2+^ sources. Similar results were observed in PIGBOS-knockdown cells (Fig. S3A).

**Figure 3.**
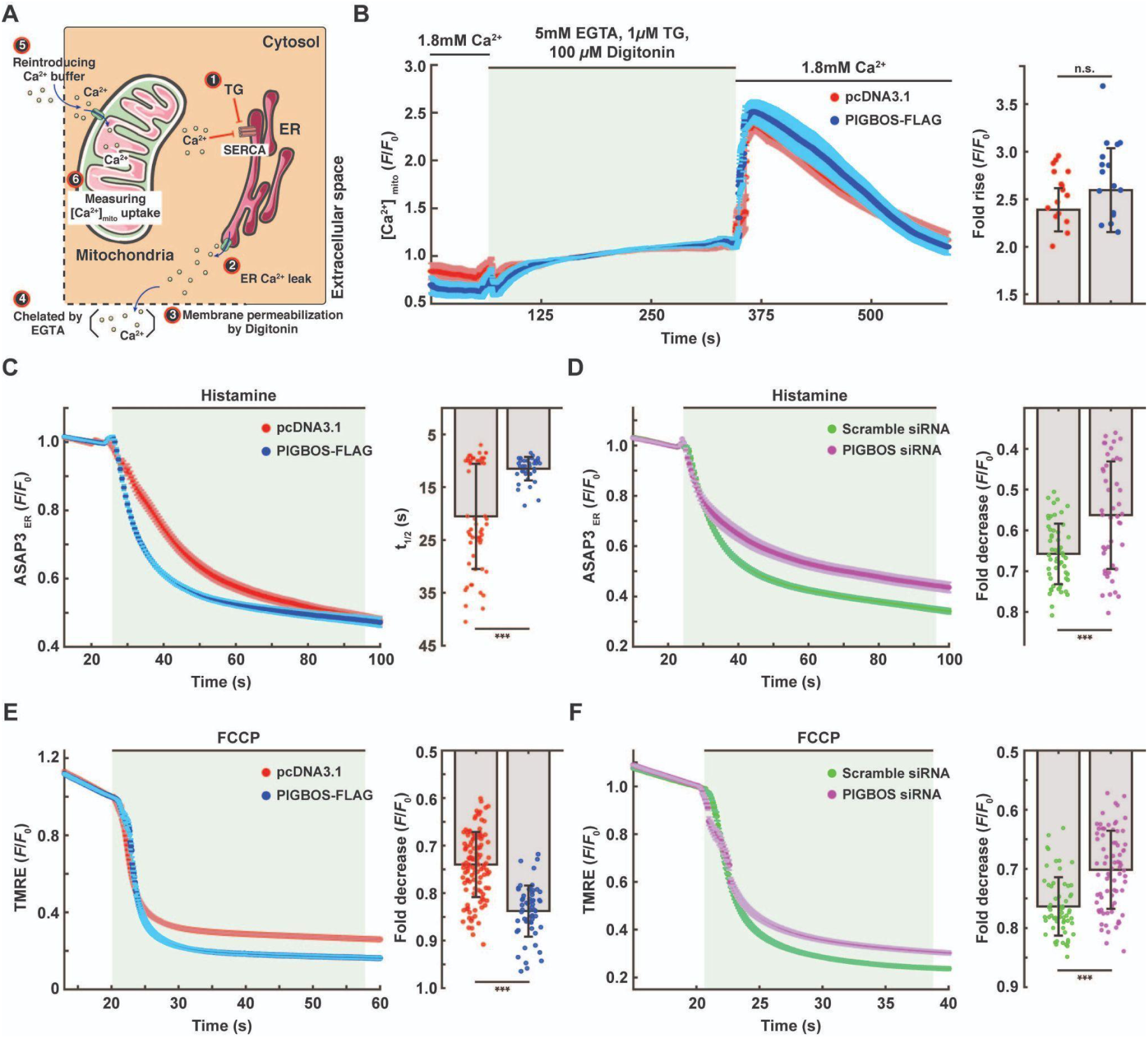
PIGBOS modulates ER–mitochondria Ca^2+^ transfer and alters organellar membrane potentials. (A) Schematic of the experimental protocol for assessing ER-independent mitochondrial Ca^2+^ uptake in digitonin-permeabilized cells. Sequential steps (1–6): (1) thapsigargin (TG) inhibits SERCA, triggering ER Ca^2+^ release through leak channels (2); (3) plasma membrane permeabilization allows EGTA to chelate residual cytosolic Ca^2+^ (4); (5) reintroduction of Ca^2+^-containing buffer enables mitochondrial uptake; (6) mitochondrial Ca²⁺ rise is measured using the targeted indicator GCEPIA1-2mt. (B) Mitochondrial Ca^2+^ uptake from the extracellular buffer is unchanged upon PIGBOS overexpression (*n* = 16 cells). (C) ER membrane depolarizes more rapidly after histamine stimulation in PIGBOS-overexpressing cells. *t*_1/2_ indicates the time to 50% reduction in fluorescence. Right, quantification of ER membrane depolarization (PIGBOS-overexpressing, *n* = 60; control, *n* = 77). (D) Histamine-induced ER membrane depolarization is attenuated in PIGBOS-depleted cells (*n* = 45) relative to scrambled controls (*n* = 56). (E) FCCP-induced mitochondrial depolarization monitored by TMRE fluorescence. PIGBOS-overexpressing cells (*n* = 59) show greater FCCP-induced decay than controls (*n* = 117), indicating higher basal mitochondrial membrane potential. (F) PIGBOS knockdown decreases mitochondrial depolarization compared with scrambled siRNA controls (PIGBOS-depleted, *n* = 76; control, *n* = 60), suggesting reduced mitochondrial membrane potential. Data are mean ± SD. n.s., not significant; ***P < 0.001 (unpaired t test).

### PIGBOS modulates ER and mitochondrial membrane potentials

Given the effects of PIGBOS on ER-mitochondria Ca^2+^ flux, we next examined whether PIGBOS influences the ER and mitochondrial membrane potentials. Using the ER-targeted voltage indicator ASAP3_ER_, we measured changes in ER membrane potential (ΔΨ_ER_) following histamine stimulation. PIGBOS overexpression led to a faster decrease in fluorescence, indicating accelerated ER depolarization. In control cells, a 50% fluorescence reduction occurred within ∼20 s, whereas in PIGBOS-overexpressing cells, it occurred within ∼12 s (Fig. 3C). Conversely, PIGBOS knockdown cells exhibited smaller depolarization responses, consistent with reduced [Ca^2+^]_ER_ release (Fig. 3D). This implies that higher release of ER Ca^2+^ in PIGBOS-overexpressing cells creates more ER depolarization, and vice-versa during PIGBOS knockdown.

Mitochondrial membrane potential (ΔΨ_m_) was measured using TMRE. Application of FCCP caused a loss of ΔΨ_m_, reflected by a decrease in TMRE fluorescence. PIGBOS-overexpressing cells exhibited a larger fluorescence drop, suggesting an elevated basal ΔΨ_m_ - a hallmark of enhanced mitochondrial function (Fig. 3E). In contrast, PIGBOS knockdown and KO cells showed a smaller decrease in TMRE fluorescence, indicative of lower basal ΔΨ_m_, implying that these cells suffer from compromised mitochondrial health. (Fig. 3F, Fig. S3B).

### PIGBOS influences store-operated Ca^2+^ entry

Since we observed that manipulation of PIGBOS levels alters ER Ca^2+^ stores and their homeostasis, we next examined whether PIGBOS influences store-operated Ca^2+^ entry (SOCE), which is activated upon depletion of ER Ca^2+^ stores. To assess SOCE, the SERCA pump was inhibited with TG in Ca^2+^-free extracellular solution to deplete ER Ca^2+^ stores, followed by re-addition of 1.8 mM Ca^2+^ to trigger SOCE. In PIGBOS-overexpressing cells, TG elicited a smaller increase in cytosolic Ca²⁺ under Ca^2+^-free conditions, indicating reduced ER Ca^2+^ leak/depletion, but resulted in a significantly enhanced SOCE response upon Ca^2+^ re-addition (Fig. 4A). In contrast, PIGBOS knockdown cells showed no significant change in TG-induced ER Ca^2+^ release, yet exhibited markedly reduced SOCE amplitudes (Fig. 4B). Notably, PIGBOS KO cells displayed increased TG-induced ER Ca^2+^ leak, which was followed by a diminished SOCE response (Fig. S4A).

**Figure 4.**
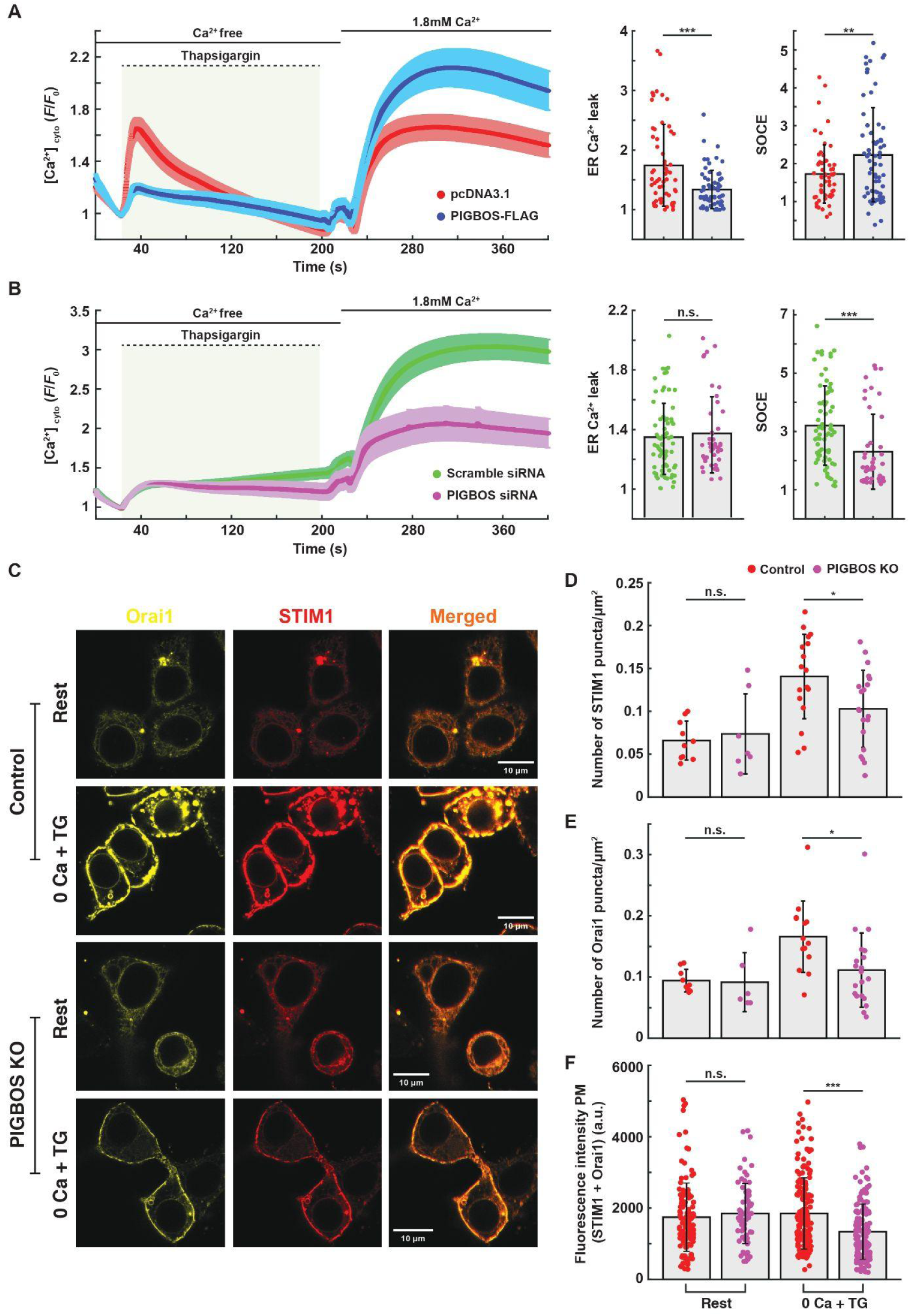
PIGBOS modulates store-operated Ca^2+^ entry (SOCE). (A) TG-induced ER Ca^2+^ leak and SOCE recordings in HEK293T cells show reduced ER Ca^2+^ leak and enhanced SOCE in PIGBOS-overexpressing cells (*n* = 66) compared with mock controls (*n* = 58). (B) PIGBOS knockdown (*n* = 44) leaves ER Ca^2+^ leak unchanged but markedly reduces SOCE relative to control cells (*n* = 71). (C) Confocal images showing STIM1 and Orai1 puncta formation following TG-induced ER Ca^2+^ store depletion in control and PIGBOS KO cells. (D and E) Quantification of STIM1 (*n* = 7-18 cells) and Orai1 (*n* = 6-18 cells) puncta before and after TG treatment. (F) Combined plasma membrane fluorescence intensity of STIM1 and Orai1 before and after TG stimulation. Each data point represents a manually defined ROI along the plasma membrane. Data represent mean ± SD. n.s., not significant; *P < 0.05, **P < 0.01, ***P < 0.001 (unpaired t test).

One possible explanation for the altered SOCE is an alteration in STIM1-Orai1 interactions. To test this, YFP-tagged Orai1 and mCherry-tagged STIM1 were expressed in PIGBOS KO and control cells. The number of STIM1 and Orai1 puncta, reflecting their interaction at ER–plasma membrane junctions, was quantified before and after TG treatment. Notably, upon TG stimulation, PIGBOS KO cells exhibited significantly fewer STIM1 and Orai1 puncta compared with control cells (Fig. 4C–E). Consistently, the combined plasma membrane fluorescence intensity of STIM1 and Orai1 was reduced in PIGBOS KO cells (Fig. 4F), indicating weakened STIM1-Orai1 coupling in the absence of PIGBOS. To determine whether PIGBOS also affects STIM1 and Orai1 expression, we performed western blot analysis. Both STIM1 and Orai1 protein levels increased upon PIGBOS overexpression and decreased following PIGBOS knockdown or knockout (Fig. S4B). Collectively, these findings suggest that PIGBOS positively regulates both the expression and interaction of STIM1 and Orai1, thereby modulating SOCE.

### PIGBOS-CLCC1 interaction is crucial for Ca^2+^ signaling

Chu *et al.* previously demonstrated that PIGBOS interacts with CLCC1 via its C-terminal region (20). To assess the functional significance of this interaction in terms of mediating ER-mitochondria Ca^2+^ homeostasis, we generated a C-terminal deletion mutant lacking residues 30–54 (ΔC-PIGBOS) (Fig. S5A), cloned into pICherryNeo backbone. Consistent with previous reports, ΔC-PIGBOS localized normally to mitochondria (Fig. S5B) but failed to enhance histamine-induced cytosolic or mitochondrial Ca^2+^ transients compared with full-length PIGBOS (Fig. 5A-B). We next tested whether ΔC-PIGBOS could rescue the Ca^2+^ signaling defects in PIGBOS KO cells. While full-length PIGBOS restored the histamine-induced [Ca^2+^]_cyto_ increase in PIGBOS KO cells, ΔC-PIGBOS did not (Fig. 5C), indicating that the C-terminal region of PIGBOS is essential for its effects on ER Ca^2+^ release.

**Figure 5.**
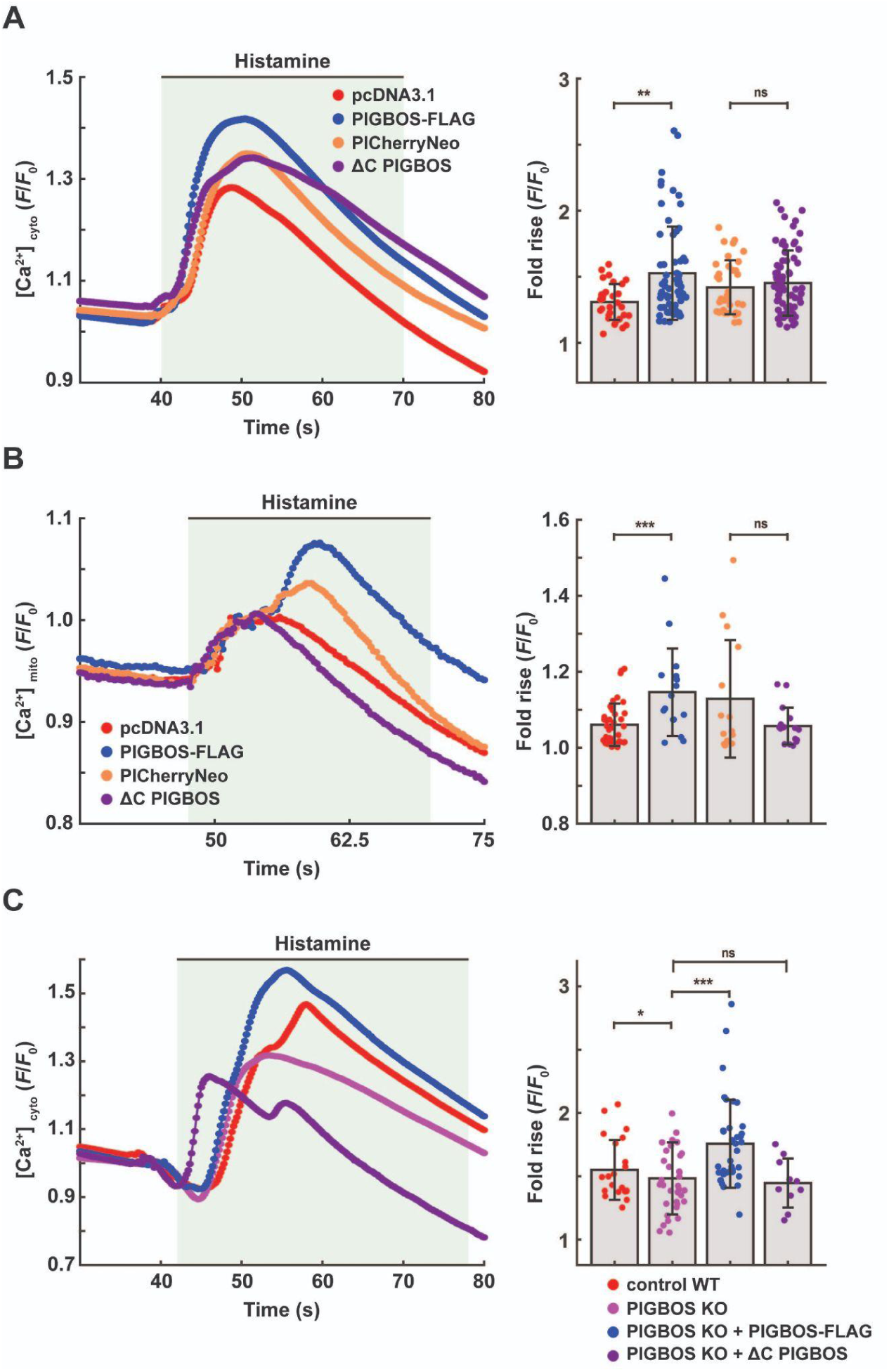
The C-terminus of PIGBOS is essential for modulating ER Ca^2+^ release and subsequent mitochondrial uptake. (A) Expression of full-length PIGBOS (WT), but not C-terminally truncated PIGBOS (ΔC PIGBOS), significantly enhanced histamine-induced cytosolic Ca^2+^ elevations (*n* = 32, 64, 35, and 73 cells for pcDNA3.1, PICherryNeo, PIGBOS-FLAG, and ΔC PIGBOS, respectively). (B) WT PIGBOS expression also augmented histamine-evoked mitochondrial Ca^2+^ responses, whereas ΔC PIGBOS expression attenuated them (*n* = 40, 16, 15, and 17 cells, respectively). (C) Re-expression of WT PIGBOS in PIGBOS-KO HEK293T cells restored the histamine-induced cytosolic Ca^2+^ rise (third bar from left), whereas ΔC PIGBOS failed to rescue the phenotype (fourth bar). Untransfected PIGBOS-KO cells exhibited significantly reduced Ca^2+^ transients (second bar) compared to cells with endogenous PIGBOS (first bar, control WT). *n* = 52, 49, 40, and 12 cells for control WT, KO, KO + PIGBOS-FLAG, and KO + ΔC PIGBOS, respectively. Data represent mean ± SD. *P < 0.05, **P < 0.01 (unpaired t test).

To further probe the role of CLCC1 in cellular Ca^2+^ homeostasis, we reduced its expression by ∼40% using shRNA (Fig. 6A). CLCC1 knockdown phenocopied PIGBOS depletion, resulting in reduced histamine-triggered [Ca^2+^]_ER_ release (Fig. 6B), lower basal [Ca^2+^]_ER_ levels (Fig. 6C), diminished ER depolarization (Fig. 6D), and decreased mitochondrial Ca^2+^ uptake (Fig. 6E). Together, these results demonstrate that the PIGBOS-CLCC1 interaction is essential for coupling ER and mitochondrial Ca^2+^ signaling.

**Figure 6.**
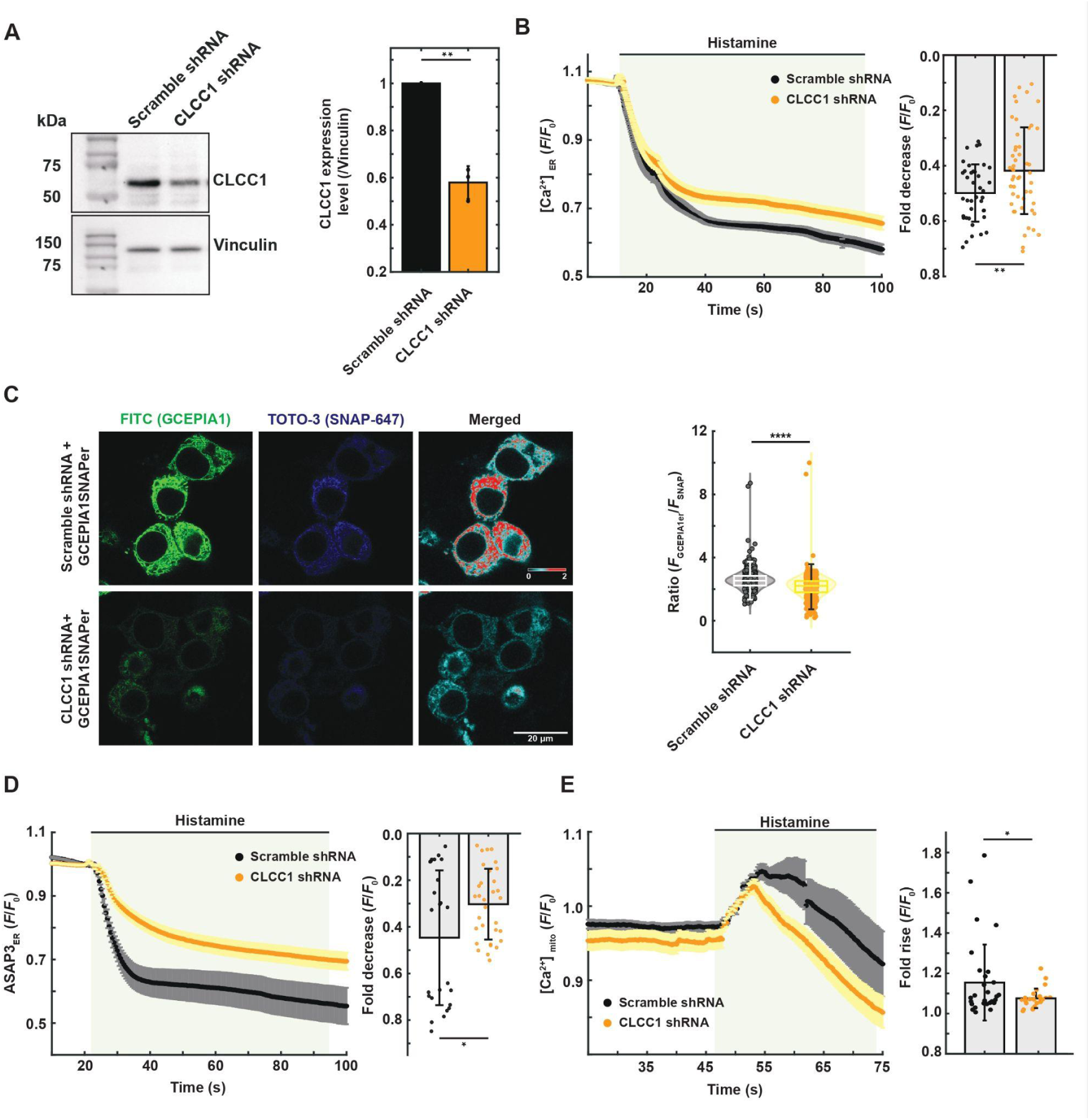
CLCC1 depletion alters histamine-induced ER Ca^2+^ release, ER potential change and mitochondrial Ca^2+^ uptake. (A) Representative western blot showing reduced CLCC1 protein expression relative to vinculin following shRNA-mediated knockdown. The accompanying bar graph shows the densitometric quantification of CLCC1 protein levels (*N* = 3). (B) ER Ca^2+^ release following histamine stimulation. CLCC1 knockdown decreases histamine-induced ER Ca^2+^ release compared with control cells. The bar graph summarizes data from 39 control and 50 knockdown cells. (C) Basal ER Ca^2+^ levels imaged with GCEPIA1-SNAP_ER_. Representative confocal images of scramble shRNA-transfected control and CLCC1-knockdown cells are shown. The violin plot summarizes quantified fluorescence intensity from 387-419 cells of control and knock-down cells, respectively, indicating a significant reduction in basal ER Ca^2+^ content in CLCC1-depleted cells. (D) ER membrane potential changes in response to histamine. CLCC1-deficient cells (*n* = 32) show a smaller histamine-induced ER voltage change than scramble controls (*n* = 25), as summarized in the bar graph. (E) Mitochondrial Ca^2+^ dynamics following histamine treatment. Histamine-induced mitochondrial Ca^2+^ increase is significantly lower in CLCC1-knockdown cells (*n* = 25) compared with control cells (*n* = 31). Data are presented as mean ± SD. Statistical significance was determined using an unpaired t-test: n.s., not significant; *P < 0.05; **P < 0.01; ****P < 0.0001.

### PIGBOS is associated with the Ca^2+^ signaling network

To assess whether PIGBOS is a functional component of the cellular Ca^2+^ signaling network, we performed a STRING-based protein–protein interaction analysis. PIGBOS was directly linked to CLCC1, which in turn was predicted to associate with key Ca^2+^ signaling components, including STIM1, SERCA, Orai1, and IP_3_R (Fig. S6A). To test these network predictions, we quantified mRNA expression of representative Ca^2+^ signaling genes by RT-qPCR. PIGBOS overexpression increased CLCC1, IP_3_R, and SERCA, whereas PIGBOS knockdown had opposite effects (Fig. S6B), indicating that PIGBOS influences the transcriptional landscape of the Ca^2+^ signaling pathway. PIGBOS KO also showed a similar trend to knockdown cells (Fig. S6C).

### PIGBOS modulates mitochondrial metabolism and redox balance

Cytosolic and mitochondrial Ca^2+^ levels critically regulate mitochondrial metabolism and oxygen consumption rate (OCR), with reduced mitochondrial Ca^2+^ ([Ca^2+^]_mito_) or [Ca^2+^]_cyto_ often associated with impaired respiration (23–26). Given the role of PIGBOS in Ca^2+^ homeostasis, we assessed mitochondrial function using Seahorse metabolic flux analysis. PIGBOS overexpression enhanced basal respiration and ATP production without significantly altering proton leak or spare respiratory capacity (Fig. 7A–B). In contrast, PIGBOS knockdown or knockout decreased all respiratory parameters (Fig. 7C–D; Fig. S7A–B) and increased mitochondrial ROS levels, indicating elevated oxidative stress (Fig. 7E).

**Figure 7.**
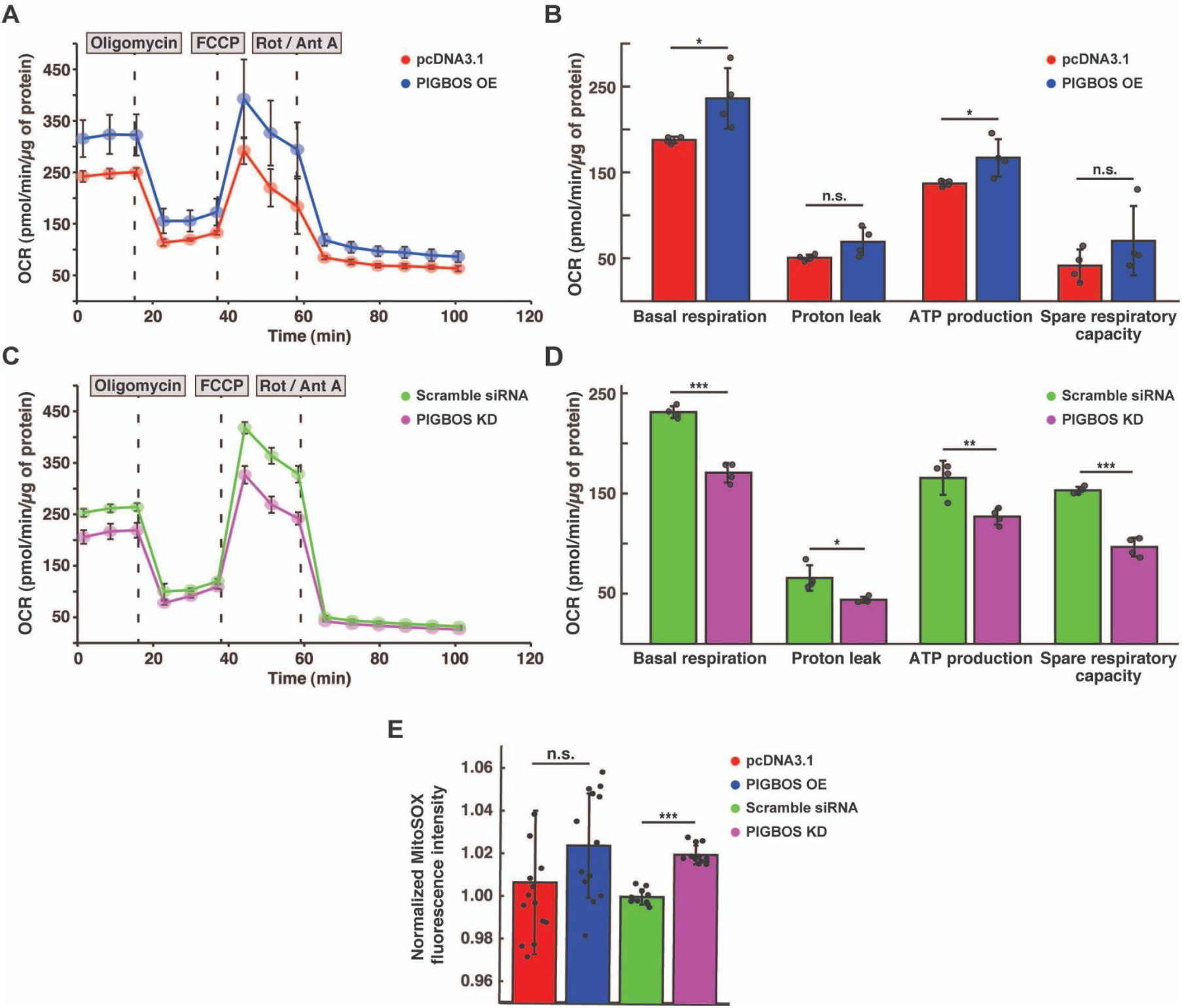
PIGBOS modulates mitochondrial metabolism and reactive oxygen species (ROS) levels. (A, B) Overexpression of PIGBOS in HEK293T cells increases basal respiration and ATP production compared to empty vector controls (*N* = 3). (C, D) Conversely, PIGBOS knockdown reduces basal respiration, ATP production and spare respiratory capacity relative to scramble controls (*N* = 3). (E) Mitochondrial ROS levels are elevated in PIGBOS knockdown cells compared with controls, whereas PIGBOS overexpression does not significantly alter ROS levels (*N* = 13). Data are presented as mean ± SD. n.s., not significant; *P < 0.05, **P < 0.01, ***P < 0.001 (unpaired t-test).

## Discussion

Recent studies have identified numerous functional microproteins, typically under 100 amino acids in length, encoded by small open reading frames (sORFs) (27). These microproteins regulate essential cellular processes such as ion transport, oxidative phosphorylation, and stress responses (28, 29). Several localize to the sarco/endoplasmic reticulum (SR/ER) and mitochondria, where they control key functions including Ca^2+^ homeostasis and ATP production (30–33). In the SR, the microprotein DWORF activates SERCA (34), whereas Phospholamban (35), Sarcolipin (36,37), Myoregulin (38), and Endoregulin (39) inhibit it. Together, these regulators fine-tune cytosolic Ca^2+^ clearance, [Ca^2+^]_ER_ levels, and downstream processes such as muscle contraction and apoptosis (40–42). Mitochondria localized microproteins like ASAP (ATP synthase–associated peptide) (43), BRAWNIN (44, 45), Humanin (46, 47), Mitoregulin (48), and MOTS-c (49) regulate several core aspects of mitochondrial metabolism (50).

In 2019, Chu *et al.* (20) discovered PIGBOS, a microprotein localized to the mitochondrial outer membrane that interacts with CLCC1 and modulates ER stress. However, the implications of PIGBOS-CLCC1 interaction in the context of ER-mitochondria signaling were not explored. It is possible that PIGBOS acts through its interacting partner, CLCC1. Since ER stress can result from dysregulated [Ca^2+^]_ER_ homeostasis and CLCC1 contributes to [Ca^2+^]_ER_ balance (22), we hypothesized that PIGBOS may play a key role in the cellular Ca^2+^ signaling network. Previously, we reported that PIGBOS overexpression in HEK293T cells enhanced histamine/ionomycin-induced intracellular Ca^2+^ rise and mitochondrial Ca^2+^ uptake (51). In the present study, we provide an in-depth analysis showing that the PIGBOS-CLCC1 interaction governs intracellular Ca^2+^ dynamics, regulating Ca^2+^ transfer from the ER to mitochondria and thereby influencing metabolic coupling. The ER stress–regulatory role of PIGBOS possibly arises from its control over cellular Ca^2+^ signaling, particularly through maintaining ER Ca^2+^ homeostasis.

PIGBOS overexpression elevates basal [Ca^2+^]_ER_ and enhances histamine-induced Ca^2+^ release, whereas its knockdown or knockout reduces both parameters (Fig. 1 and 2). Mechanistically, PIGBOS sustains ER ionic balance by interacting with CLCC1, which provides counter-ion conductance required for optimum Ca^2+^ transport (22). Deletion of the CLCC1-binding C-terminal region of PIGBOS abolishes its effect on Ca^2+^ signaling (Fig. 5A-B), and ΔC-PIGBOS fails to rescue the signaling defects in PIGBOS-null cells (Fig. 5C). Similarly, CLCC1 knockdown decreases basal [Ca^2+^]_ER_, histamine-induced Ca^2+^ release, and mitochondrial Ca^2+^ uptake, mirroring the PIGBOS knockdown phenotype (Fig. 6F, 6H). These findings indicate that PIGBOS positively regulates CLCC1 function and acts through it to modulate [Ca^2+^]_ER_ dynamics.

Although the precise mechanism of CLCC1-mediated Ca^2+^ regulation requires further investigation, previous studies showed that CLCC1 depletion lowers basal [Ca^2+^]_ER_ and Ca^2+^ release while increasing steady-state [Cl⁻]_ER_ and [K^+^]_ER_ (22). During [Ca^2+^]_ER_ release, Cl⁻ efflux and compensatory K^+^ influx maintain charge balance and osmolarity; depletion of CLCC1 may disturb this equilibrium, increasing luminal osmolarity and causing ER swelling, which subsequently hampers proper Ca^2+^ signaling (22). Consequently, PIGBOS knockdown likely produces similar effects by reducing CLCC1 activity. Additionally, PIGBOS overexpression may enhance SERCA function, given that CLCC1 is predicted to interact directly with SERCA (Fig. S6A) and SERCA levels rise upon PIGBOS expression (Fig. S6B), thereby contributing to elevated basal [Ca^2+^]_ER_.

We found that PIGBOS enhances ER-to-mitochondria Ca^2+^ transfer in a CLCC1-dependent manner without affecting Ca^2+^ uptake from extracellular sources. Deletion of the CLCC1-interacting domain of PIGBOS abolished this effect, confirming the necessity of this specific interaction. It is well established that MAMs play a critical role in facilitating IP_3_R-mediated Ca^2+^ transfer from the ER to mitochondria. Interestingly, previous studies showed that PIGBOS is not an integral component of MAMs and that its deletion does not alter ER-mitochondria tethering sites (20), a change typically observed upon loss of canonical MAM-tethering proteins. Our findings suggest that the PIGBOS–CLCC1 complex may modulate local Ca^2+^ microdomains at MAMs, fine-tuning ER ionic concentrations to influence IP_3_R-mediated Ca^2+^ release and subsequent mitochondrial uptake.

PIGBOS plays a broader role than regulating ER–mitochondria Ca^2+^ signaling; it also influences SOCE, a key pathway for Ca^2+^ influx in most cells. Cells overexpressing PIGBOS show higher SOCE, whereas its depletion produces the opposite effect. Interestingly, PIGBOS overexpression increases the levels of STIM1 and Orai1, the principal components of the SOCE machinery, while PIGBOS knockdown and knockout reduce their levels. Further, the coupling of STIM1-Orai1 interaction is lower in PIGBOS knockout cells, validating the lower SOCE response. Because PIGBOS enhances functional ER–mitochondria Ca^2+^ coupling and strengthens IP₃R-mediated Ca^2+^ release, it may create local ER Ca^2+^ depletion zones that activate STIM and open ORAI channels more efficiently.

At the systems level, PIGBOS overexpression and depletion altered the expression of CLCC1 and several genes associated with the core Ca^2+^ signaling pathway, indicating that PIGBOS links local ionic changes to global transcriptional adaptation, possibly through Ca^2+^-responsive transcription factors. Functional analysis of mitochondria showed that PIGBOS overexpression increases respiration and ATP production, whereas its depletion reduces oxidative phosphorylation and elevates ROS levels. These observations position PIGBOS as a microprotein that connects Ca^2+^ signaling to mitochondrial energy metabolism and redox balance through its interaction with CLCC1.

Although CLCC1 has been implicated in several diseases, such as ALS (22) and autosomal recessive retinitis pigmentosa (52), PIGBOS has not yet been associated with any disease, likely due to its recent discovery. Considering its critical role in Ca^2+^ signaling, ER stress, and mitochondrial function, it is reasonable to predict that future studies will reveal its involvement in diseases such as Parkinson’s disease, Alzheimer’s disease, or cancer, where Ca^2+^ signaling, the unfolded protein response (UPR), and mitochondrial functions are disrupted. Therefore, PIGBOS may emerge as a potential drug target for multiple diseases in the near future. In summary (Fig. 8), we demonstrate that PIGBOS coordinates plasma membrane-ER-mitochondria Ca^2+^ communication, coupling ionic balance to bioenergetic output. This work expands the paradigm that small mitochondrial proteins can exert broad regulatory influence on inter-organelle signaling and cellular stress adaptation.

**Figure 8.**
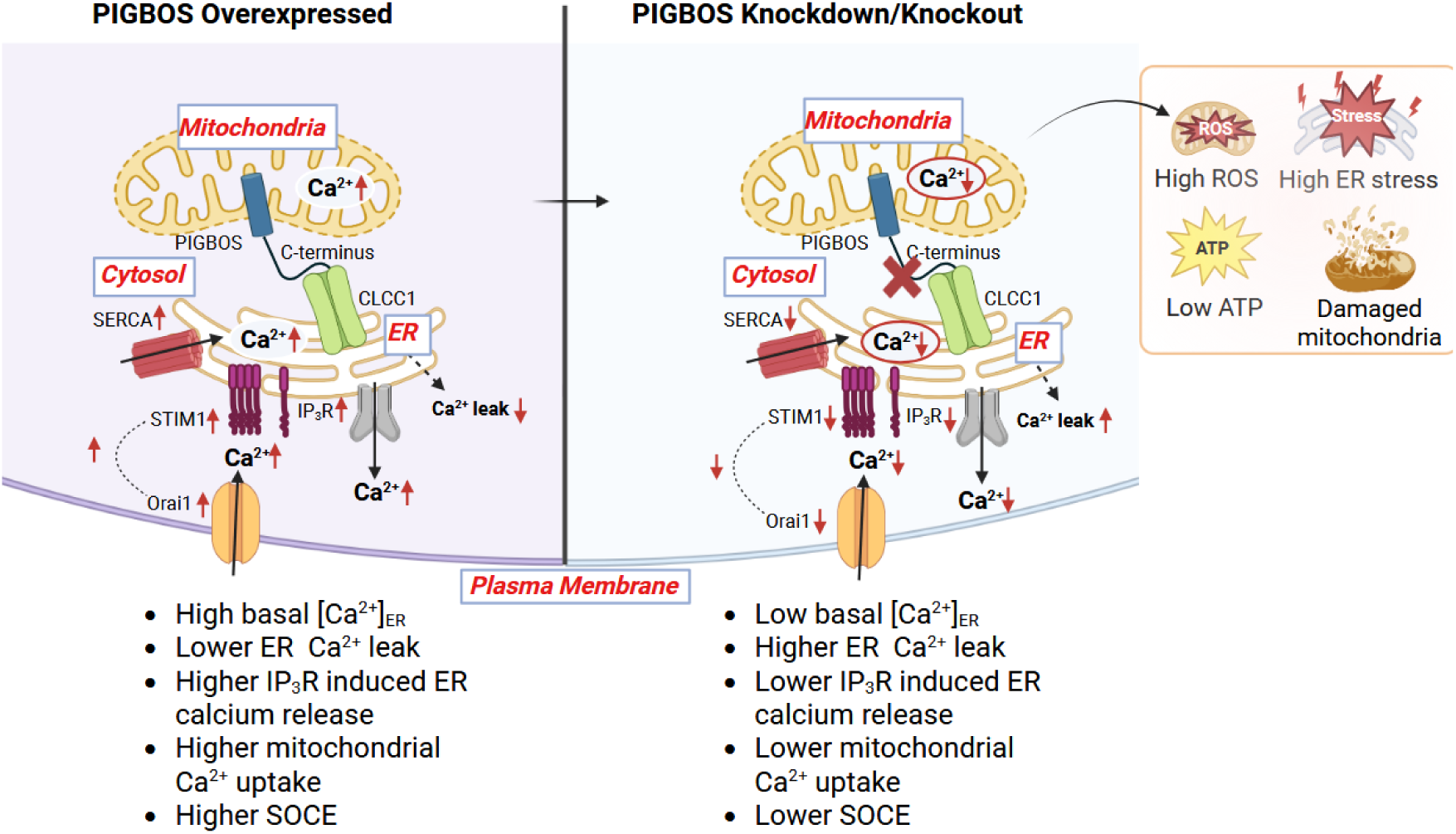
Schematic representation of how PIGBOS affects cellular Ca^2+^ homeostasis involving ER-mitochondria-plasma membrane. Figure 8. Model summarizing how PIGBOS coordinates ER-mitochondria-plasma membrane Ca^2+^ signalling. PIGBOS engages CLCC1 via its C terminus to augment IP_3_R-dependent agonist-evoked ER Ca^2+^ release, and STIM1-Orai1 coupling to potentiate SOCE, thereby amplifying cytosolic and mitochondrial Ca^2+^ transients and supporting bioenergetic and redox homeostasis. Conversely, PIGBOS loss attenuates IP_3_R, STIM1, and Orai1 function, blunts ER-to-mitochondria Ca^2+^ transfer, and promotes ROS accumulation, ATP depletion, and mitochondrial injury (figure created with BioRender).

## Limitations

Although we presented a detailed study demonstrating the role of PIGBOS in inter-organelle Ca^2+^ signaling and SOCE, our work has several limitations. All experiments were performed in HEK293T cells, which raises concerns about the generalizability of our findings to other cell types. Since our primary goal was to delineate the molecular mechanism of PIGBOS in intracellular Ca^2+^ flux rather than to characterize cell-type-specific physiological outputs, we believe that our mechanistic insights reflect core features of Ca^2+^ signaling networks. The mechanism we propose, in which the PIGBOS-CLCC1 interaction regulates [Ca^2+^]_ER_ homeostasis and facilitates IP_3_R-mediated Ca^2+^ transfer to mitochondria, is the best fit with our data and existing literature. However, because the precise role of CLCC1 in [Ca^2+^]_ER_ regulation is not fully understood, further studies are needed to validate this model using the CLCC1 mutant that lacks PIGBOS binding capacity. We identified PIGBOS as a positive modulator of CLCC1, but this requires confirmation by examining how PIGBOS influences CLCC1 channel activity in lipid bilayer. Finally, *in vivo* validation is essential to establish the translational relevance of our findings. Future studies using PIGBOS KO animal models will be crucial to assess the physiological consequences of impaired PIGBOS-mediated Ca^2+^ homeostasis.

## Materials and Methods

### Cell culture and transfection

HEK293T cells were procured from the National Centre for Cell Science (NCCS), India. Cells were maintained in Dulbecco’s Modified Eagle Medium (DMEM) supplemented with 10% heat-inactivated fetal bovine serum (FBS) and 1x antibiotic–antimycotic solution (Gibco #154240-062). Cells were cultured under standard conditions at 37°C in a cell culture incubator maintaining 5% CO_2_ levels and passaged when approximately 80% confluency was reached. For imaging experiments, cells were seeded at equal densities on 35 mm dishes containing glass coverslips.

For transfection, cells were grown to 50–60% confluency and transfected using JetPrime Transfection Reagent (Polyplus #10100051) according to the manufacturer’s instructions. A total of 2–3 µg of plasmid DNA was used per transfection, with equal amounts of each construct when multiple plasmids were co-transfected. For western blotting and RNA isolation, cells were transfected under identical conditions after plating at equal densities in 60 mm dishes. Culture medium was replaced 5 h post-transfection.

In overexpression experiments, control cells were transfected with the empty pcDNA3.1 vector to equalize total DNA content with groups overexpressing PIGBOS-FLAG. For knockdown assays, a scrambled siRNA served as the negative control, and a PIGBOS-specific siRNA (Dharmacon #N-192024-01-0002) was used for targeted silencing. The scrambled siRNA (MISSION siRNA Universal Negative Control #1, SIC001) was purchased from Sigma-Aldrich.

### Generation of PIGBOS knockout HEK293T cells

CRISPR–Cas9 constructs were generated by cloning a single-guide RNA (sgRNA) sequence reported by Chu et al. (20), which targets exon 2 of the PIGBOS1 locus, into the PX458 vector backbone (Addgene #48138). HEK293T cells were transfected with the resulting CRISPR–Cas9 plasmids, and GFP-positive cells were isolated 48 h post-transfection by fluorescence-activated cell sorting (FACS) into 96-well plates. Single-cell–derived PIGBOS KO clones were expanded and screened by western blot analysis to confirm loss of PIGBOS expression.

### Plasmids

pcDNA3.1(+) was used as the empty vector control for overexpression experiments. The PIGBOS–FLAG plasmid was kindly provided by Dr. A. Saghatelian (Salk Institute). Additional constructs included pCMV G-CEPIA1_ER_ (Addgene #58215) and pCMV CEPIA-2mt (Addgene #58218) from Dr. M. Iino; GCEPIA1–SNAP_ER_ (Addgene #172659) from Dr. P. Xu; pGP-CMV-jGCaMP7s (Addgene #104463) from Dr. D. Kim and the GENIE Project; and pCAG-ASAP3_ER_ (Addgene #195356) from Dr. B. Bloodgood. Orai-YFP and STIM1-mCherry plasmids were kindly provided by Dr. Shmuel Muallem (National Institute of Dental and Craniofacial Research, USA). Full-length PIGBOS and a C-terminal deletion (ΔC) mutant were cloned into pICherryNeo (Addgene #52119). For knockdown experiments, CLCC1 shRNA (Sigma-Aldrich #TRCN0000257146) and a scrambled shRNA control (Addgene #1864) were used.

### Mitochondria isolation and western blotting

Mitochondria were isolated as described (53) with minor modifications. Cells were lysed in isolation buffer (75 mM sucrose, 225 mM mannitol, 5 mM HEPES pH 7.4, 1 mM EGTA) with PMSF and protease inhibitors (Sigma #P8340), vortexed 30 min, and centrifuged (1,300 x g, 5 min x 3) to remove cellular debris. The supernatant was centrifuged (13,000 x g, 25 min) to pellet mitochondria, which were then lysed in RIPA buffer. Protein concentration was determined by the Lowry method (54). Samples were resolved on 12% Tricine gels, transferred to PVDF membranes, blocked in 5% BSA, and probed with antibodies against PIGBOS (Abgenex #ABP121, 1:500), CLCC1 (Thermo Fisher #A305-668A-T, 1:1,000), STIM1 (CST #D88E10, 1:1,000), Orai1 (Novus #NBP1-85463, 1:1,000), VDAC (CST #4866, 1:1,000), and vinculin (CST #4650, 1:5,000), followed by HRP-conjugated secondary antibodies: anti-rabbit (CST #7074, 1:5,000) and anti-mouse (CST #7076, 1:5,000) and Clarity ECL detection (Bio-Rad #1705060).

### Confocal microscopy

Cells were seeded on 18-mm square glass coverslips (Bluestar), transfected the following day, and, at 48 h post-transfection, labeled with 20 nM MitoTracker Red CMXRos (Thermo Fisher #M7512) for 30 min at 37°C, 5% CO_2_. Cells were fixed in 4% paraformaldehyde (Himedia #TCL119), permeabilized with 0.1% Triton X-100, blocked in 5% BSA/PBS for 1 h, and stained with anti-FLAG primary antibody (CST #14793S, 1:100, overnight at 4°C) followed by Alexa Fluor secondary antibody (Invitrogen #A11034, 1 h at room temperature); nuclei were counterstained with DAPI and coverslips were mounted in DPX (SRL #88417). Images were acquired using a 60 x oil-immersion objective on an Olympus FluoView 3000 confocal microscope.

For STIM1–Orai1 puncta analysis, cells seeded on 35-mm glass-bottom dishes were co-transfected with STIM1-mCherry and Orai1-YFP, 24 h later were treated with 2 μM thapsigargin (Sigma #T9033) for 5 min or left untreated. They were then imaged pre- and post-treatment using a 60 x oil-immersion objective (excitation: YFP 514 nm; mCherry 561 nm) on an Olympus FluoView 3000 confocal microscope.

### Ca^2+^ imaging

HEK293T cells were cultured on 12-mm glass coverslips (Bluestar) and transfected with organelle-specific Ca^2+^ sensor plasmids, the indicated PIGBOS constructs, and siRNAs according to the respective experimental conditions. 48 h post-transfection, cells were washed with Hank’s balanced salt solution (HBSS; 140 mM NaCl, 5.6 mM KCl, 3.6 mM NaHCO_3_, 1.8 mM CaCl_2_, 1 mM MgCl_2_, 5.6 mM glucose, 10 mM HEPES, pH 7.4) and transferred to a perfusion chamber (Warner Instruments) continuously supplied with HBSS. Fluorescence imaging was performed using an Olympus IX83 inverted microscope equipped with an sCMOS camera (ORCA-Flash4.0, Hamamatsu Photonics), appropriate dichroic mirrors and filter sets (Chroma Technology Corp.), and an LED light source (CoolLED pE-340fura). Exposure time and light intensity were maintained constant across all recordings. Time-lapse images were acquired at 200–250 ms intervals following agonist or test compound addition.

Cytosolic Ca^2+^ dynamics were monitored using the genetically encoded sensor jGCaMP7s (55). Ca^2+^ release from the ER was triggered by histamine (100 µM). For steady-state [Ca^2+^]_ER_ measurements, HEK293T cells expressing GCEPIA1-SNAP_ER_ (56) were plated on 35-mm glass-bottom dishes, washed once with HBSS, and incubated with SNAP-Cell 647-SiR (New England Biolabs, #S9106) for 45 min at 37°C.

Following two washes, cells were imaged in HBSS using a 60 x oil immersion objective on an Olympus FluoView 3000 confocal microscope (excitation: 640 nm, emission: 661 nm). CEPIA1 fluorescence intensity was normalized to SNAP fluorescence in each cell to account for expression variability, as described previously (56).

Mitochondrial Ca^2+^ levels were measured using the mitochondria-targeted reporter CEPIA2-mt. For mitochondrial Ca^2+^ uptake assays in permeabilized cells, HEK293T cells co-transfected with GCEPIA2-mt (57), were washed with HBSS and transferred to a perfusion chamber. Images were recorded continuously for 5 min. During the first minute, cells were perfused with HBSS, followed by a Ca^2+^-free, high-K^+^ permeabilization buffer (125 mM KCl, 5 mM NaCl, 2 mM K_2_HPO_4_, 1 mM MgCl_2_, 20 mM HEPES, pH 7.2, supplemented with 5 mM glutamate, 100 µM digitonin, 5 mM EGTA, and 1 µM thapsigargin). Thapsigargin depleted ER Ca^2+^ stores, and EGTA chelated residual Ca^2+^ in the digitonin-permeabilized cells. After 2 min, HBSS containing 1.8 mM Ca^2+^ was reintroduced, eliciting a rapid increase in GCEPIA2-mt fluorescence indicative of mitochondrial Ca^2+^ uptake from the extracellular medium.

Store-operated Ca²⁺ entry (SOCE) was assessed by first depleting ER Ca^2+^ stores using a Ca^2+^-free buffer (140 mM NaCl, 5.6 mM KCl, 3.6 mM NaHCO_3_, 1 mM MgCl_2_, 5 mM EGTA, 5.6 mM glucose, 10 mM HEPES, pH 7.4) containing 0.5 µM thapsigargin. After ∼3.5 min of ER depletion and passive Ca^2+^ leak, HBSS supplemented with 1.8 mM Ca^2+^ was reintroduced to induce SOCE.

### Measurement of ER voltage

ER membrane potential dynamics were monitored using the ER-targeted voltage sensor ASAP3_ER_ (58). Upon histamine stimulation, a decrease in fluorescence intensity indicated ER membrane depolarization. Images were captured every 250 ms, and fluorescence traces were analyzed offline using ImageJ and custom MATLAB scripts.

### Mitochondrial membrane potential and ROS measurement

Mitochondrial membrane potential was assessed using tetramethylrhodamine ethyl ester (TMRE). Cells on coverslips were incubated in HBSS containing 200 nM TMRE for 5 min at room temperature in the dark, followed by two washes with HBSS. Time-lapse imaging was performed (excitation: 562 nm; emission: 641 nm), and 1 µM FCCP was added after 20 s to induce mitochondrial depolarization.

Mitochondrial ROS was measured using MitoSOX™ Red (Thermo Fisher #M36008). Cells were incubated with 2.5 µM MitoSOX in HBSS for 30 min at 37°C with 5% CO_2_, washed twice with HBSS, and imaged on an Olympus IX83 microscope using an mCherry filter set.

### Image analysis and quantification

For Ca^2+^ imaging experiments, individual cells were manually delineated as regions of interest (ROIs) in ImageJ following background subtraction. Fluorescence intensity values across time were extracted using the Time Series Analyser plugin and processed in MATLAB (R2020b, MathWorks). The baseline fluorescence immediately preceding agonist application was defined as F_0_. Ca^2+^ transients were identified using the cusum algorithm, and fluorescence changes were normalized as F/F_0_. The half-time (*t*_1/2_) was computed as the time to reach 50% of maximum or minimum fold change relative to F_0_.

STIM1–Orai1 puncta were quantified after background subtraction using the *Analyze Particles* function in ImageJ (3–100 pixels, circularity 0.1–1.0). Data were expressed as puncta per µm^2^. For plasma membrane fluorescence analysis, ROIs were manually defined, and fluorescence intensities from YFP and mCherry channels were summed to assess STIM1-Orai1 colocalization before and after thapsigargin treatment. Figures were generated in MATLAB and finalized in Adobe Illustrator (Adobe Inc.).

### mRNA expression analysis

Cells were washed with ice-cold PBS, and total RNA was extracted using TRIzol reagent (Takara #9108). The aqueous RNA phase was isolated following chloroform extraction, precipitated with isopropanol, and washed with 70% ethanol. cDNA was synthesized using the cDNA Conversion Kit (Bio-Rad #170-8891). Quantitative real-time PCR was performed using 200 nM of each primer (listed in Supplementary Table 1). Gene expression was normalized to β-actin, which served as an internal reference.

### Mitochondrial respiration analysis

Mitochondrial function in HEK293T cells was measured using the Seahorse XF96 Analyzer (Agilent Technologies) according to the manufacturer’s protocol. Approximately 5 × 10^4^ cells per well were seeded onto Seahorse XF96 cell culture plates and equilibrated in assay medium (DMEM supplemented with 200 mM glutamine, 100 mM pyruvate, and 1 M glucose). The Seahorse XF Cell Mito Stress Test was performed using sequential injections of oligomycin (1.5 µM), FCCP (1 µM), and a mixture of rotenone and antimycin A (0.5 µM each). Oxygen consumption rate (OCR) was recorded and normalized to protein content for intersample comparison.

### Statistical analysis

All experiments were repeated at least 3 times unless otherwise stated. Statistical comparisons between two groups were performed using unpaired Student’s t tests. Data are presented as mean ± SD. Statistical significance is denoted as P < 0.05 (*), P < (**), P < 0.001 (***), and P < 0.0001 (****).

## Declaration of AI Use

ChatGPT (GPT-3.5) was used to refine language and readability. The authors reviewed all content and assumed full responsibility for the final version of this manuscript.

## Acknowledgments

The work was funded by the intramural fund provided by the Department of Biotechnology, IIT Madras. S.A. is supported by the Prime Minister’s Research Fellowship (PMRF), Ministry of Education, Government of India. The authors thank the BIO-SAIF facility, Department of Biotechnology, IIT Madras, for assistance with confocal imaging and flow cytometry. We acknowledge Dr. Pratyay Sengupta for guidance with calcium imaging data analysis, and Sambhavi Pattnaik, Satrajit Das, and Roshni Raj for critical manuscript review. We also thank the contributors who provided plasmids used in this study.

## Author Contributions

SA and AKB conceptualized the study and designed the research; SA performed all experiments and analyzed the data; SA and AKB wrote the paper; AKB secured funding.

## Competing Interests

The authors declare that they have no competing interests.

## Notes

### Competing Interest Statement

The authors have declared no competing interest.

## References

1. M. Brini, T. Cali, D. Ottolini, & E. Carafoli, Neuronal calcium signaling: function and dysfunction. Cell. Mol. Life Sci 71, 2787–2814 (2014).

2. J. Humeau, J.M.B. Pedro, I. Vitale, L. Nuñez, C. Villalobos, G. Kroemer, L. Senovilla, Calcium signaling and cell cycle: Progression or death. Cell Calcium 70, 3–15 (2018).

3. A. E. Carafoli, Calcium signaling: a tale for all seasons. Proc. Natl. Acad. Sci. U. S. 99, 1115–22 (2002).

3. S. Luan, C. Wang, Calcium Signaling Mechanisms Across Kingdoms. Annu. Rev. Cell Dev. Biol. 37, 311–340 (2021).

4. S. Zheng et al., Calcium homeostasis and cancer: insights from endoplasmic reticulum-centered organelle communications. Trends Cell Biol. 33, 312–323 (2023).

5. E. Neher, T. Sakaba, Multiple Roles of Calcium Ions in the Regulation of Neurotransmitter Release. Neuron 59, 861–872 (2008)

6. M.D. Bootman, G. Bultynck, Fundamentals of Cellular Calcium Signaling: A Primer. Cold Spring Harb. Perspect. Biol. 12, a038802 (2020).

7. R. Bagur et al., Intracellular Ca^2+^ Sensing: Its Role in Calcium Homeostasis and Signaling. Mol. Cell 66, 780–788 (2017).

8. M.J. Berridge, The endoplasmic reticulum: a multifunctional signaling organelle. Cell Calcium 32, 235–249 (2002).

9. D. Billups, B. Billups, R. A. J. Challiss, & S. R. Nahorski, Modulation of Gq-protein-coupled inositol trisphosphate and Ca^2+^ signaling by the membrane potential. J. Neurosci. 26, 9983–9995 (2006).

10. J.P. Decuypere, G. Monaco, L. Missiaen, H. De Smedt, J.B. Parys, G. Bultynck, IP_3_ Receptors, Mitochondria, and Ca^2+^ Signaling: Implications for Aging. J. Aging Res. 2011, 920178 (2011).

11. J. Krebs, L.B. Agellon, M. Michalak, Ca^2+^ homeostasis and endoplasmic reticulum (ER) stress: An integrated view of calcium signaling. Biochem. Biophys. Res. Commun. 460, 114–121 (2015).

12. Daverkausen-Fischer, Lea et al., Regulation of calcium homeostasis and flux between the endoplasmic reticulum and the cytosol. J. Biol Chem. 298, 102061 (2022).

13. K. Princen et al., Pharmacological modulation of septins restores calcium homeostasis and is neuroprotective in models of Alzheimer’s disease. Science 384, eadd6260 (2024)

14. G. Morciano, A. Rimessi, S. Patergnani, V.A.M. Vitto, A. Danese, A. Kahsay, L. Palumbo, M. Bonora, M. Wieckowski, C. Giorgi, P. Pinton, Calcium dysregulation in heart diseases: Targeting calcium channels to achieve a correct calcium homeostasis. Pharmacol. Res. 177, 1043–6618 (2022).

15. F. Hadi, M. Mortaja, & Z. Hadi, Calcium (Ca^2+^) hemostasis, mitochondria, autophagy, and mitophagy contribute to Alzheimer’s disease as early moderators. Cell Biochem. Funct. 42, e4085 (2024).

16. N. Kumari, N. Pullaguri, R.S. Narayan and A. Bajaj, V. Sahu, & K.K.R. Ealla, Dysregulation of calcium homeostasis in cancer and its role in chemoresistance. Cancer Drug Resist. 7, 11 (2024).

17. A. Raffaello, C. Mammucari, G. Gherardi, & R. Rizzuto, Calcium at the Center of Cell Signaling: Interplay between Endoplasmic Reticulum, Mitochondria, and Lysosomes. Trends Biochem. Sci. 41, 1035–1049 (2016).

18. W.B. Zhao, & R. Sheng, The correlation between mitochondria-associated endoplasmic reticulum membranes (MAMs) and Ca^2+^ transport in the pathogenesis of diseases. Acta Pharmacol. Sin. 46, 271–291 (2025).

19. Q. Chu, T.F. Martinez, S.W. Novak et al., Regulation of the ER stress response by a mitochondrial microprotein. Nat. Commun. 10, 4883 (2019).

20. Y. Jia, T.J. Jucius, S. A. Cook, & S.L. Ackerman, Loss of Clcc1 Results in ER Stress, Misfolded Protein Accumulation, and Neurodegeneration. J. Neurosci. 35, 3001–3009 (2015).

21. L. Guo, Q. Mao, J. He et al., Disruption of ER ion homeostasis maintained by an ER anion channel CLCC1 contributes to ALS-like pathologies. Cell Res. 33, 497–515 (2023).

22. A. Rossi, P. Pizzo, & R. Filadi, Calcium, mitochondria and cell metabolism: A functional triangle in bioenergetics. Biochim. Biophys. Acta Mol. Cell Res. 1866, 1068–1078 (2019).

24. A. L. Piao, Y.H. Fang, M. Fisher, R.B. Hamanaka, A. Ousta, R. Wu, G.M. Mutlu, A. J. Garcia 3rd, S.L. Archer, W.W. Sharp, Dynamin-related protein 1 is a critical regulator of mitochondrial calcium homeostasis during myocardial ischemia/reperfusion injury. FASEB J. 38, e23379 (2024).

23. E.A. Vilas-Boas, J.V. Cabral-Costa, V.M. Ramos, C.C. Caldeira da Silva, & A.J. Kowaltowski, Goldilocks calcium concentrations and the regulation of oxidative phosphorylation: Too much, too little, or just right. J. Biol. Chem. 299, 102904 (2023).

24. G. Gherardi, L. Nogara, S. Ciciliot, G.P. Fadini, B. Blaauw, P. Braghetta, P. Bonaldo, D. De Stefani, R. Rizzuto, C. Mammucari, Loss of mitochondrial calcium uniporter rewires skeletal muscle metabolism and substrate preference. Cell Death Differ. 26, 362–381 (2019).

25. J.J. Mohsen, A.A. Martel, & S.A. Slavoff, Microproteins-Discovery, structure, and function. Proteomics 23, 23–24 e2100211 (2023).

26. K.R. Hassel, O. Brito-Estrada, & C.A. Makarewich, Microproteins: Overlooked regulators of physiology and disease. iScience 26, 106781 (2023).

27. K. Geering, FXYD proteins: new regulators of Na-K-ATPase. Am. J. Physiol. Renal Physiol. 290, F241–50 (2006).

28. T.M. Coughlin, & C.A. Makarewich, Emerging roles for microproteins as critical regulators of endoplasmic reticulum function and cellular homeostasis. Semin Cell Dev. Biol. 170, 103608 (2025).

29. M.L. Kamradt & C.A. Makarewich, Mitochondrial microproteins: Critical regulators of protein import, energy production, stress response pathways, and programmed cell death. Am. J. Physiol. Cell Physiol. 325, C807–C816 (2023).

30. K. Yen, B. Miller, H. Kumagai, A. Silverstein & P. Cohen, Mitochondrial-derived microproteins: from discovery to function. Trends Genet. 41, 132–145 (2025).

31. N. Borghol, S. Yandiev & J. Courchet, Mitochondrial microproteins: Emerging regulators in neurodevelopment and neurodegeneration. Bioessays 47, e70058 (2025).

32. B.R. Nelson, C.A. Makarewich, D.M. Anderson, B. R. Winders, C.D. Troupes, F. Wu, A.L. Reese, J.R. McAnally, X. Chen, E.T. Kavalali, S.C. Cannon, S.R. Houser, R. Bassel-Duby, E.N. Olson, A peptide encoded by a transcript annotated as long noncoding RNA enhances SERCA activity in muscle. Science 351, 271–275 (2016).

33. F. Funk, A. Kronenbitter, K. Hackert, M. Oebbeke, G. Klebe, M. Barth, D. Koch, J. P. Schmitt, Phospholamban pentamerization increases sensitivity and dynamic range of cardiac relaxation. Cardiovasc. Res. 119, 1568–1582 (2023).

34. G. J. Babu, P. Bhupathy, V. Timofeyev, N.N. Petrashevskaya, P.J. Reiser, N. Chiamvimonvat, M. Periasamy, Ablation of sarcolipin enhances sarcoplasmic reticulum calcium transport and atrial contractility. Proc. Natl. Acad. Sci. U. S. A. 104, 17867–17872 (2007).

35. C. Montigny, D.L. Huang, Beswick, V., et al., Sarcolipin alters SERCA1a interdomain communication by impairing binding of both calcium and ATP. Sci. Rep. 11, 1641 (2021).

36. D. M. Anderson, K.M. Anderson, C. Chang, C.A. Makarewich, B.R. Nelson, J.R. McAnally, P. Kasaragod, J.M. Shelton, J. Liou, R. Bassel-Duby, E.N. Olson, A micropeptide encoded by a putative long noncoding RNA regulates muscle performance. Cell 160, 595–606 (2015).

37. C.A. Makarewich, K.M. Anderson, J.M. Shelton, S. Bezprozvannaya, R. Bassel-Duby, E.N. Olson, Widespread control of calcium signaling by a family of SERCA-inhibiting micropeptides. Sci. Signal. 9, ra119 (2016).

38. D.R. Singh, M.P. Dalton, E.E. Cho, M.P. Pribadi, T.J. Zak, J. Šeflová, C.A. Makarewich, E.N. Olson, S.L. Robia, Newly Discovered Micropeptide Regulators of SERCA Form Oligomers but Bind to the Pump as Monomers. J. Mol. Biol. 431, 4429–4443 (2019).

39. A. Odermatt, P.E.M. Taschner, S.W. Scherer, B. Beatty, V.K. Khanna, D.R. Cornblath, V. Chaudhry, W. Yee, B. Schrank, G. Karpati, M.H. Breuning, N. Knoers, D.H. Maclennan, Characterization of the gene encoding human sarcolipin (SLN), a proteolipid associated with SERCA1: absence of structural mutations in five patients with Brody disease. Genomics 45, 541–553 (1997).

40. S. Appleby, H.M. Aitken-Buck, M.S. Holdaway, M.S. Byers, C.M. Frampton, L.N. Paton, A.M. Richards, R.R. Lamberts, C.J. Pemberton, Cardiac effects of myoregulin in ischemia-reperfusion. Peptides 174, 171156 (2024).

41. Q. Ge, D. Jia, D. Cen, Y. Qi, C. Shi, J. Li, L. Sang, L.J. Yang, J. He, A. Lin, S. Chen, L. Wang, Micropeptide ASAP encoded by LINC00467 promotes colorectal cancer progression by directly modulating ATP synthase activity. J. Clin. Investig. 131, e152911 (2021).

42. S. Zhang, B. Reljić, C. Liang et al., Mitochondrial peptide BRAWNIN is essential for vertebrate respiratory complex III assembly. Nat. Commun. 11, 1312 (2020).

43. Y. Wang, Y. Shi, W. Li, X. Han, X. Lin, D. Liu, Y. Lin, L. Shen, Knockdown of BRAWNIN minimally affect mitochondrial complex III assembly in human cells. Biochim. Biophys. Acta Mol. Cell Res. 1871, 119601 (2024).

44. M.A. Moreno Ayala, M.F. Gottardo, C.F. Zuccato, et al., Humanin promotes tumor progression in experimental triple negative breast cancer. Sci. Rep. 10, 8542 (2020).

45. C.P. Ha, T.N.M. Hua, V.T.A. Vo, et al., Humanin activates integrin αV–TGFβ axis and leads to glioblastoma progression. Cell Death Dis. 15, 454 (2024).

46. C.S. Stein, P. Jadiya, X. Zhang, J.M. McLendon, G.M. Abouassaly, N.H. Witmer, E.J. Anderson, J.W. Elrod, R.L. Boudreau, Mitoregulin: A lncRNA-encoded microprotein that supports mitochondrial supercomplexes and respiratory efficiency. Cell Rep. 23, 3710–3720.e8 (2018).

47. J.C. Reynolds, R.W. Lai, J.S.T. Woodhead, et al., MOTS-c is an exercise-induced mitochondrial-encoded regulator of age-dependent physical decline and muscle homeostasis. Nat. Commun. 12, 470 (2021).

48. C. Liang, S. Zhang, D. Robinson, M.V. Ploeg, R. Wilson, J. Nah, D. Taylor, S. Beh, R. Lim, L. Sun, D.M. Muoio, D.A. Stroud, L. Ho, Mitochondrial microproteins link metabolic cues to respiratory chain biogenesis. Cell Rep. 40, 111204 (2022).

49. A. Seemanti, & A.K. Bera, Mitochondrial microprotein PIGBOS1 is possibly associated with cellular calcium homeostasis and neuro-protection. Indian J. Physiol. Allied Sci. 75, 44–47 (2023).

50. L. Li, X. Jiao, I. D’Atri, F. Ono, R. Nelson, C.C. Chan, N. Nakaya, Z. Ma, Y. Ma, X. Cai, L. Zhang, S. Lin, A. Hameed, B.A. Chioza, H. Hardy, G. Arno, S. Hull, M.I. Khan, J. Fasham, G.V. Harlalka, M. Michaelides, A.T. Moore, Z.H. Coban Akdemir, S. Jhangiani, J.R. Lupski, F.P.M. Cremers, R. Qamar, A. Salman, J. Chilton, J. Self, R. Ayyagari, F. Kabir, M.A. Naeem, M. Ali, J. Akram, P.A. Sieving, S. Riazuddin, E.L. Baple, S.A. Riazuddin, A.H. Crosby, J.F. Hejtmancik, Mutation in the intracellular chloride channel CLCC1 associated with autosomal recessive retinitis pigmentosa. PLoS Genet. 14, e1007504 (2018).

51. M. R. Wieckowski, C. Giorgi, M. Lebiedzinska, J. Duszynski & P. Pinton, Isolation of mitochondria-associated membranes and mitochondria from animal tissues and cells. Nat. Protoc. 4, 1582–1590 (2009).

52. O. H. Lowry, N. J. Rosebrough, A. L. Farr & R.J. Randall, Protein measurement with the folin phenol reagent. J. Biol. Chem. 193, 256–275 (1951).

53. H. Dana, Y. Sun, B. Mohar et al., High-performance calcium sensors for imaging activity in neuronal populations and microcompartments. Nat. Methods 16, 649–657 (2019).

54. C. Luo, H. Wang, Q. Liu et al., A genetically encoded ratiometric calcium sensor enables quantitative measurement of the local calcium microdomain in the endoplasmic reticulum. Biophys. Rep. 5, 31–42 (2019).

55. J. Suzuki, K. Kanemaru, K. Ishii et al., Imaging intraorganellar Ca^2+^ at subcellular resolution using CEPIA. Nat. Commun. 5, 1–13 (2014).

56. E.P. Campbell, A.A. Abushawish, L.A. Valdez, M.K. Bell, M. Haryono, P. Rangamani, B.L. Bloodgood, Electrical signals in the ER are cell type and stimulus specific with extreme spatial compartmentalization in neurons. Cell Rep. 42, 111943 (2023).

