## Supplementary Materials for "PIGBOS-CLCC1 Interaction Shapes Cellular Calcium Dynamics and Energy Metabolism"

**Supplementary Figures**

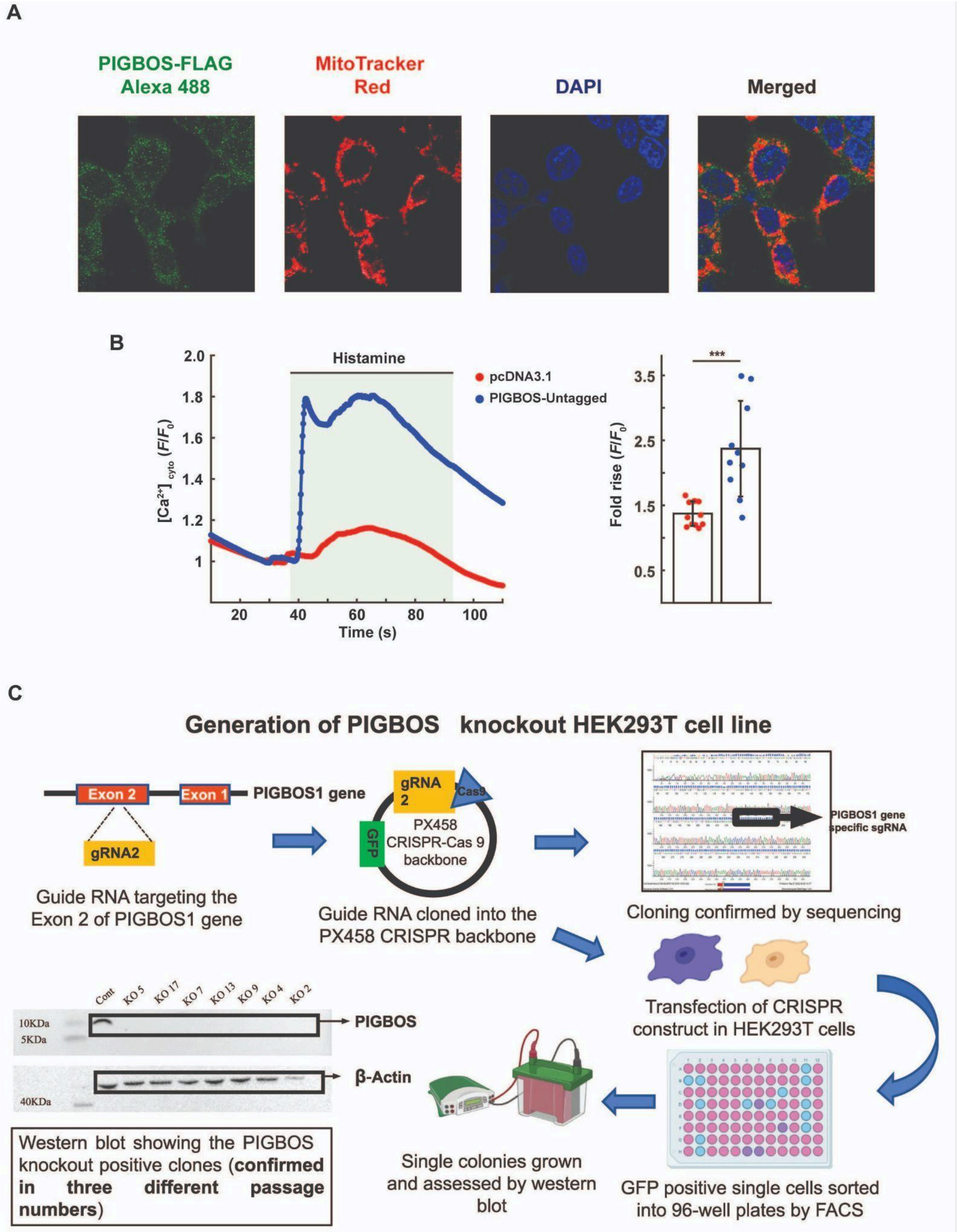


**Fig. S1.** (A) Confocal microscopy images showing mitochondrial localization of FLAG-tagged PIGBOS. PIGBOS-FLAG was detected using an Alexa Fluor 488-conjugated anti-FLAG antibody (green). Mitochondria were labelled with MitoTracker Red, and nuclei were stained with DAPI (blue). The merged image shows colocalization of PIGBOS with mitochondria, indicated by overlap of green and red signals. (B) Histamine-induced cytosolic $Ca^{2+}$ rise in HEK293T cells. Expression of untagged PIGBOS potentiated the $Ca^{2+}$ response comparable to that observed with FLAG-tagged PIGBOS (main text, Fig. 1B). Data are presented as Mean ± SD (***$P < 0.001$, unpaired t test). (C). Schematic representation of generating PIGBOS KO HEK293T cell line. PIGBOS-specific guide RNA (gRNA) targeting the exon 2 region was cloned in the CRISPR-Cas9-GFP backbone (PX458). 24 hours after transfection with the CRISPR construct containing PIGBOS gRNA, single fluorescent cells were sorted in a 96-well plate by FACS. The cells were allowed to grow for 2-3 weeks, and subsequently, samples were collected for western blot. Seven positive PIGBOS KO colonies were generated, as observed by the complete deletion of the PIGBOS band in the western blot.

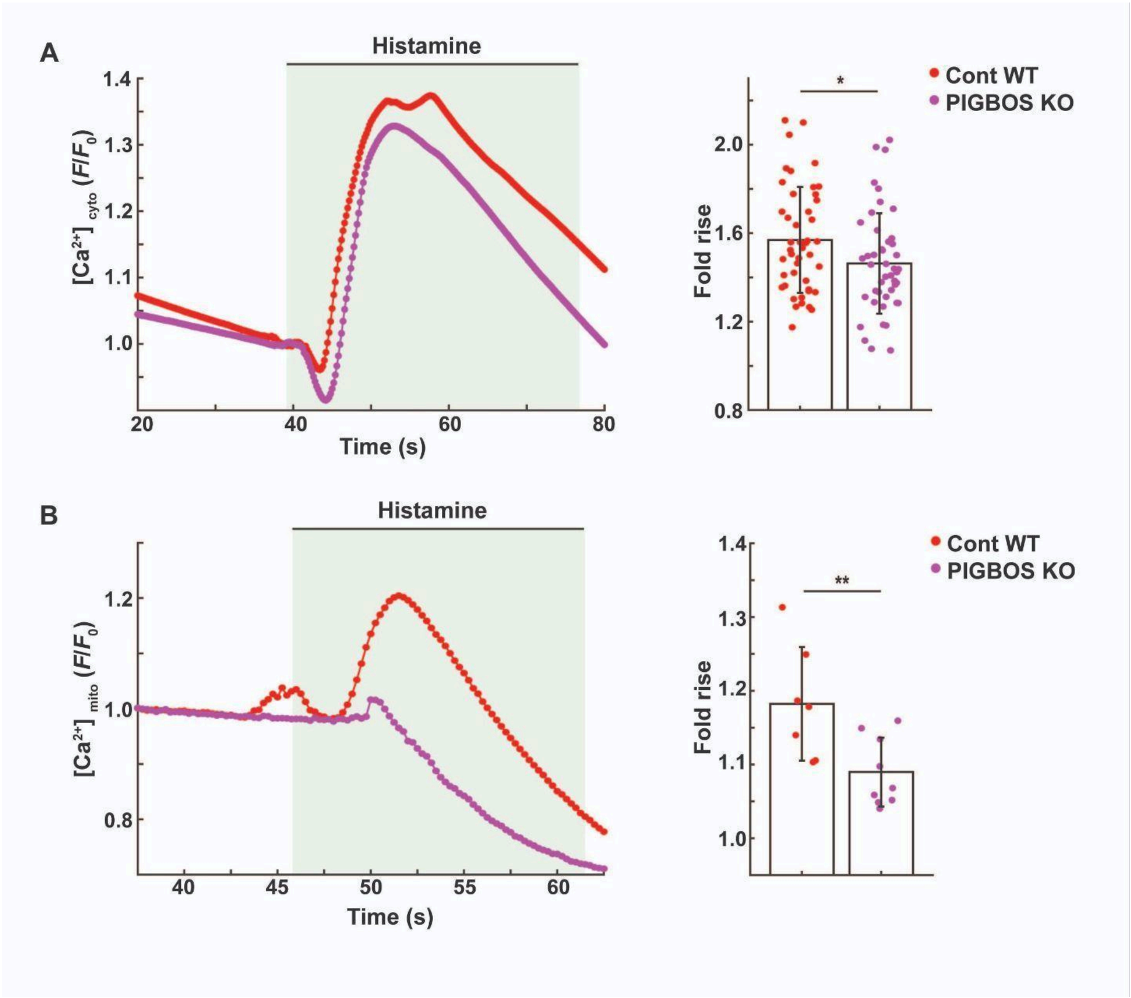


**Fig. S2.** Deletion of PIGBOS impairs histamine-evoked cytosolic $Ca^{2+}$ rise and subsequent mitochondrial $Ca^{2+}$ uptake. (A) Representative cytosolic $Ca^{2+}$ transients following histamine stimulation in control wild-type (cont WT) and PIGBOS KO HEK293T cells. The bar graph summarizes the peak cytosolic $Ca^{2+}$ responses, showing a significantly reduced $Ca^{2+}$ rise in PIGBOS1 KO cells ($n$ = 40-42 cells). Cytosolic $Ca^{2+}$ was monitored using jGCaMP7s. (B) Histamine-induced mitochondrial $Ca^{2+}$ uptake is also reduced in PIGBOS KO cells compared with control cells ($n$ = 8-9 cells), as measured with GCEPIA2-mt. Data are presented as Mean ± SD. *P < 0.05, **P < 0.01 (unpaired t test).

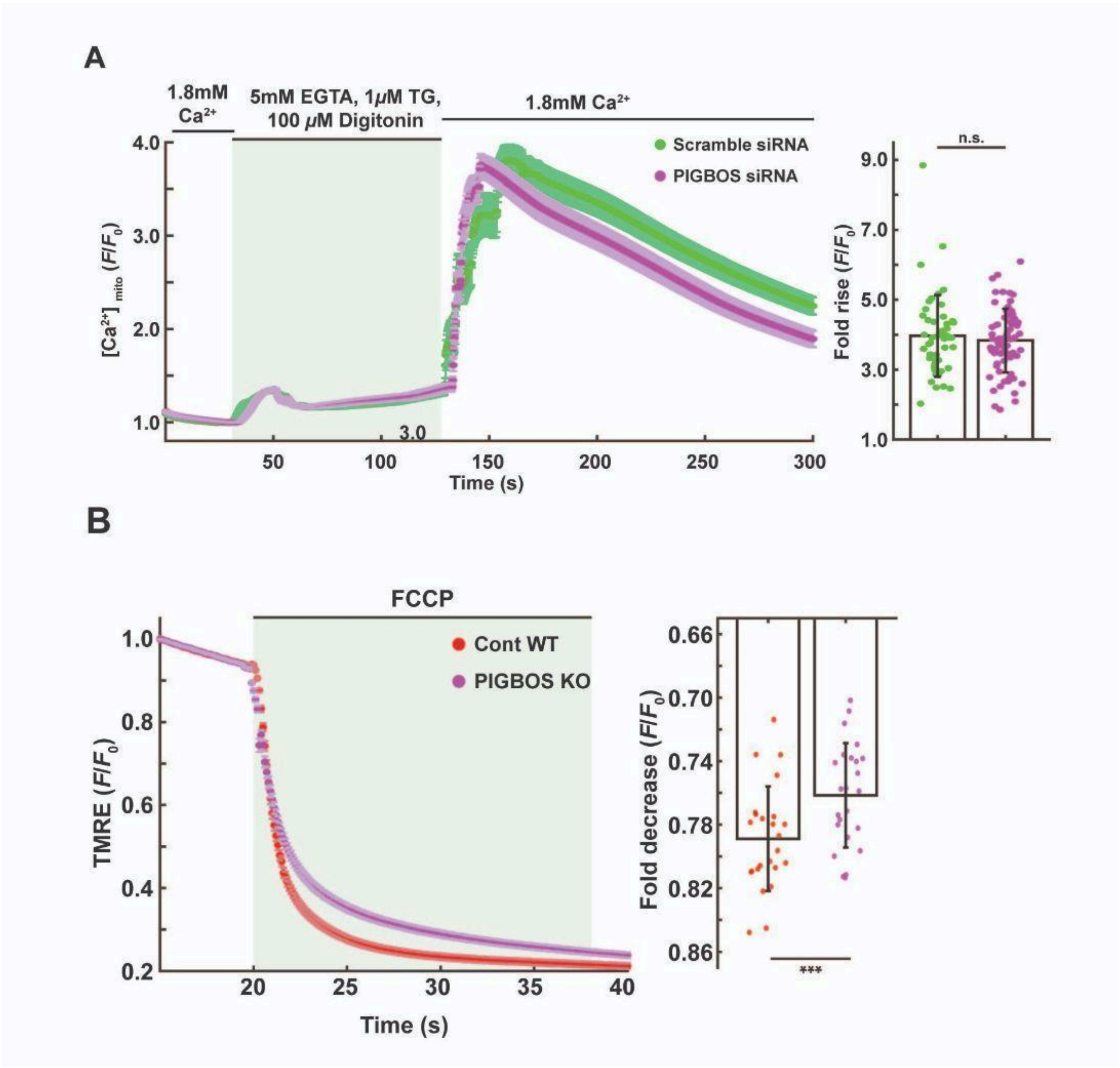


**Fig. S3.** Depletion of PIGBOS does not affect mitochondrial $Ca^{2+}$ uptake from non-ER sources but alters mitochondrial membrane potential ($\Delta\Psi_m$). (A) Representative mitochondrial $Ca^{2+}$ transients in control and siRNA-mediated PIGBOS knockdown HEK293T cells. Cells were initially perfused with $Ca^{2+}$-containing extracellular buffer, followed by $Ca^{2+}$-free buffer containing digitonin, EGTA, and thapsigargin (TG) to permeabilize the plasma membrane and deplete ER $Ca^{2+}$ stores. Subsequent readdition of extracellular $Ca^{2+}$ elicited a mitochondrial $Ca^{2+}$ rise of comparable magnitude in control and PIGBOS-depleted cells, indicating preserved mitochondrial $Ca^{2+}$ uptake from non-ER sources. The bar graph summarizes data from 45-60 cells. Mitochondrial $Ca^{2+}$ was monitored using GCEPIA-2mt. (B) Mitochondrial membrane potential ($\Delta\Psi_m$), assessed by TMRE fluorescence, was significantly reduced in PIGBOS KO cells compared with control cells. A decrease in TMRE fluorescence following FCCP treatment indicates loss of $\Delta\Psi_m$, with larger fluorescence drops reflecting higher basal $\Delta\Psi_m$. Bars represent mean ± SD from 22-24 cells. n.s., not significant; ***$P < 0.001$; (unpaired t test).

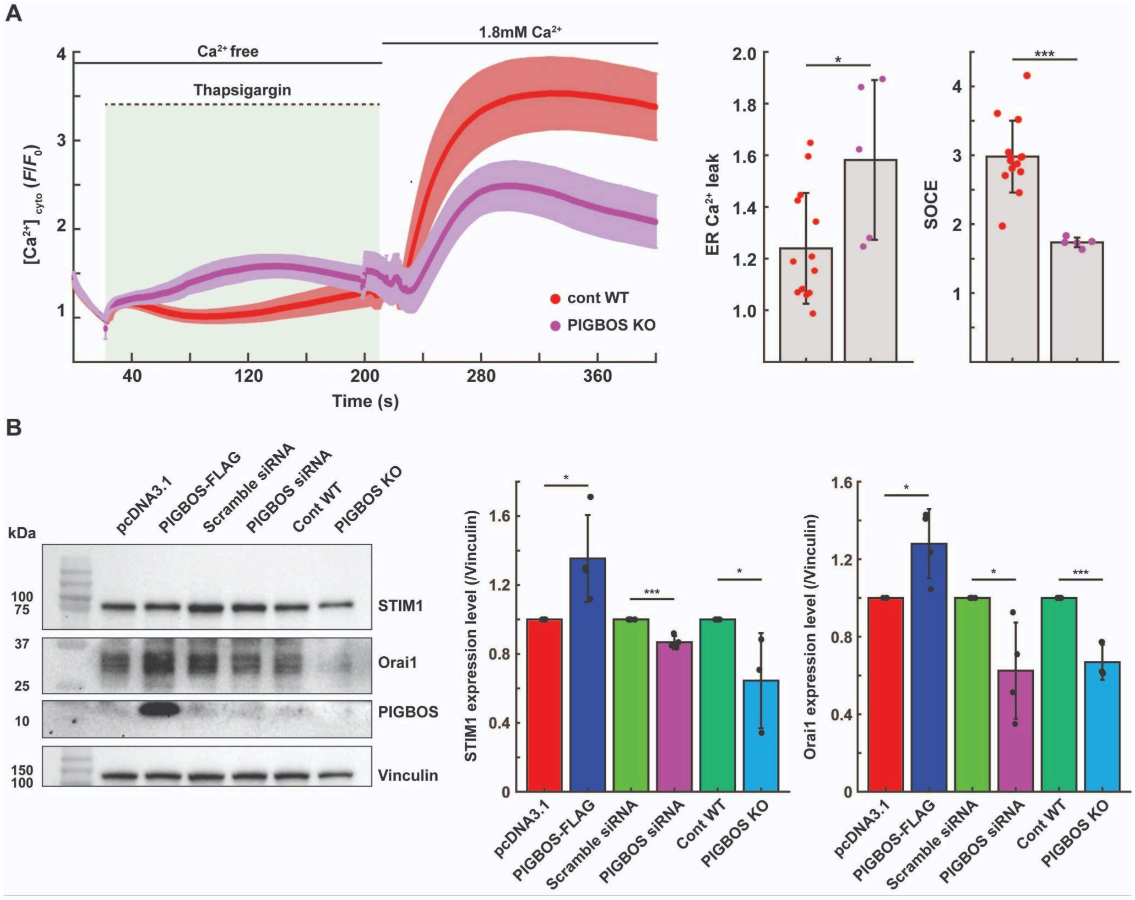


**Fig. S4.** Manipulation of PIGBOS levels alters store-operated $Ca^{2+}$ entry (SOCE) and STIM1/Orai1 expression in HEK293T cells. (A) Representative cytosolic $Ca^{2+}$ traces showing enhanced thapsigargin (TG)-induced $Ca^{2+}$ release (ER leak) but reduced SOCE (second $Ca^{2+}$ peak) in PIGBOS KO cells. cont WT: Untransfected control cells with endogenous PIGBOS. The bar graph summarizes peak $Ca^{2+}$ responses from 5-12 cells per condition. (B) Western blot analysis of STIM1 and Orai1 in HEK293T cells with PIGBOS overexpression, knockdown, or knockout. PIGBOS overexpression increases STIM1 and Orai1 protein levels, whereas PIGBOS knockdown or knockout reduces their levels. Right, quantification of fold changes in STIM1 and Orai1 normalized to loading control (vinculin) from three independent biological replicates. Data are presented as mean ± SD. *$P < 0.05$, ***$P < 0.001$ (unpaired t test).

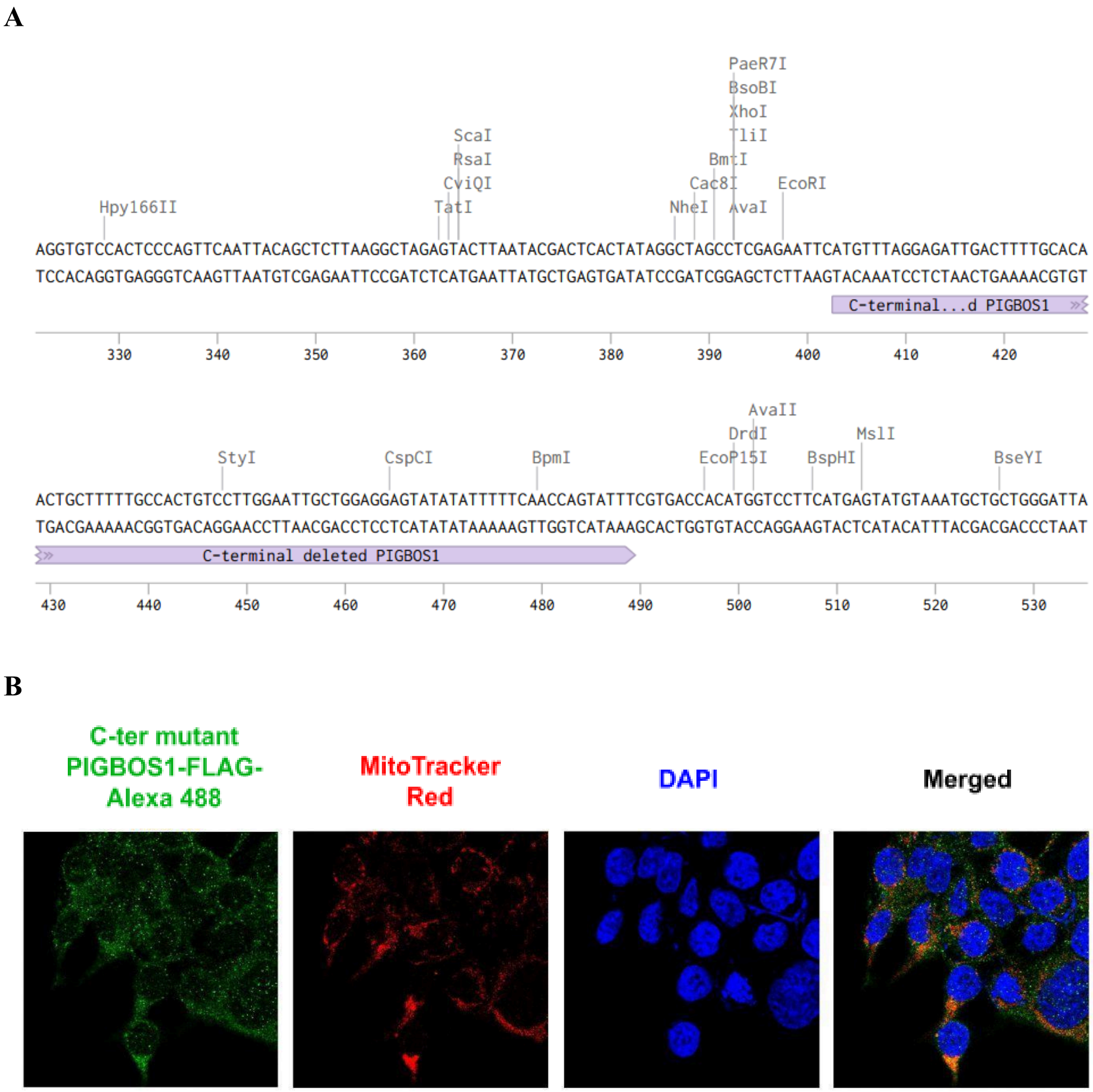


**Fig. S5.** (A) Sequencing of C-terminal deleted PIGBOS. (B) Confocal microscopy images showing mitochondrial localization of the C-terminus deleted PIGBOS (ΔC-PIGBOS) in HEK293T cells. ΔC-PIGBOS was detected using an Alexa Fluor 488-conjugated anti-FLAG antibody (green). Mitochondria were labeled with MitoTracker Red, and nuclei were stained with DAPI (blue). The merged image shows colocalization of PIGBOS with mitochondria, indicated by the overlap of green and red signals.

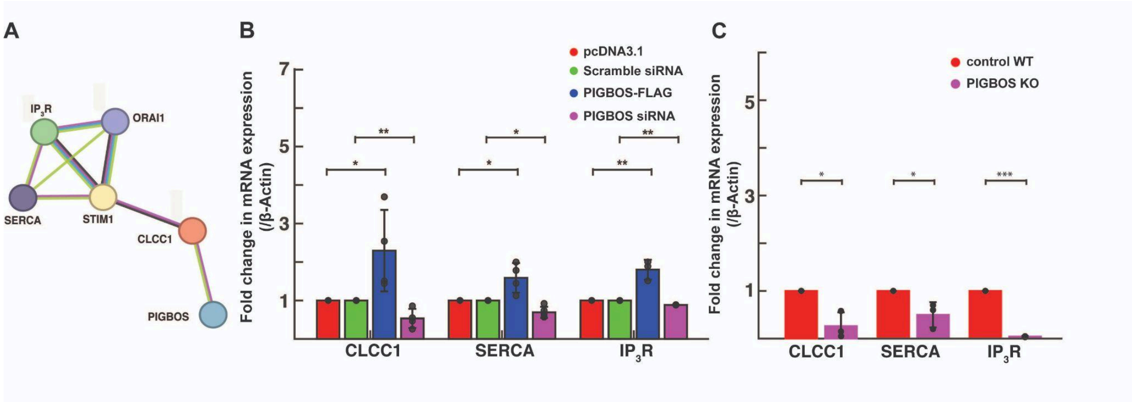


**Fig. S6.** Manipulation of PIGBOS levels alters the expression of genes associated with $Ca^{2+}$ signaling. (A) STRING network analysis showing the interaction between PIGBOS and CLCC1, which connects with multiple $Ca^{2+}$ signaling-related proteins. (B) RT-qPCR analysis demonstrating that PIGBOS overexpression upregulates CLCC1, SERCA, and IP$_3$R transcript levels, whereas PIGBOS knockdown leads to their downregulation ($n = 3$). (C) PIGBOS KO cells exhibit reduced expression of CLCC1, SERCA, and IP$_3$R, similar to PIGBOS knockdown and opposite to PIGBOS overexpression ($n = 3$). Data are presented as mean ± SD. *P < 0.05; **P < 0.01; ***P < 0.001 (unpaired t test).

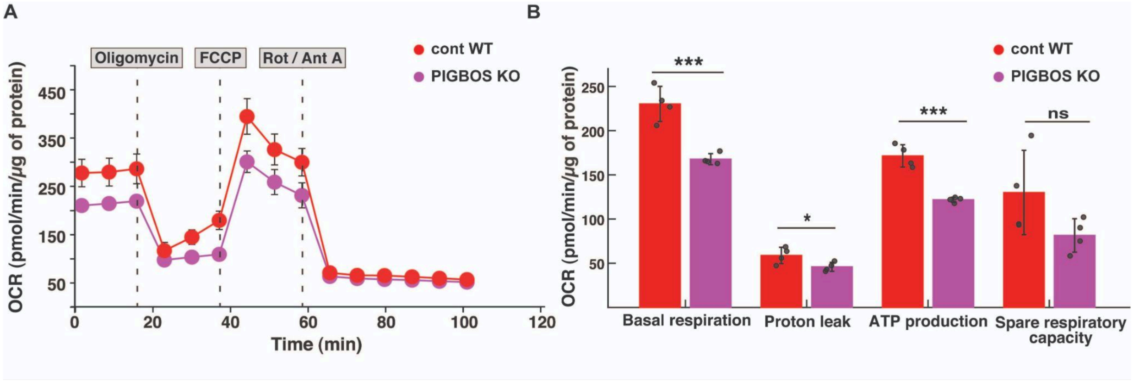


**Fig. S7.** Effect of PIGBOS deletion on mitochondrial metabolism. (A and B) Basal respiration, ATP production, and proton leak are markedly reduced in PIGBOS KO cells compared with control cells, indicating impaired mitochondrial function. Mitochondrial metabolic parameters were assessed using the Agilent Seahorse metabolic analyzer. Data are presented as mean ± SD. n.s., not significant; *P < 0.05; **P < 0.01; ***P < 0.001; (unpaired t test).

### Supplementary Table

**Supplementary Table 1: List of primers used in this study**

| **Name** | **Sequence (5’-> 3’)** |
|---|---|
| PIGBOS1_f | GGTTTCCTTTCGCCGTTTCC |
| PIGBOS1_r | CAAATCTCCGACTGTTCTCGC |
| CLCC1_f | TCCTGACTTGTCATGTGCTG |
| CLCC1_r | GAGTCAGTCAGGTGGTCTAAGT |
| SERCA_f | TCAACGAGAGTACGGGGCTG |
| SERCA_r | GTTCCAGCAAGGTTTTTCCTTCT |
| $IP_3R$_f | GAGTTTCAGCCCTCAGTGGAC |
| $IP_3R$_r | CCTTCAGGCACAGAGACCAG |
| β-Actin_f | AGCACAGAGCCTCGCCTT |
| β-Actin_r | CATCATCCATGGTGAGCTGG |

f - forward, r- reverse
